# Reversible lipid mediated pH-gating of connexin-46/50 by cryo-EM

**DOI:** 10.1101/2025.02.12.637953

**Authors:** Joshua M. Jarodsky, Janette B. Myers, Steve L. Reichow

## Abstract

Gap junctions, formed by connexin proteins, establish direct electrical and metabolic coupling between cells, enabling coordinated tissue responses. These channels universally respond to intracellular pH changes, closing under acidic conditions to limit the spread of cytotoxic signals during cellular stress, such as ischemia. Using cryo-electron microscopy (cryo-EM), we uncover insights into the structural mechanism of pH-gating in native lens connexin-46/50 (Cx46/50) gap junctions. Mild acidification drives lipid infiltration into the channel pore, displacing the N-terminal (NT) domain and stabilizing pore closure. Lipid involvement is both essential and fully reversible, with structural transitions involving an ensemble of gated-states formed through non-cooperative NT domain movement as well as minor populations of a distinct destabilized open-state. These findings provide molecular insights into pH-gating dynamics, illustrating how structural changes may regulate gap junction function under cellular stress and linking Cx46/50 dysregulation to age-related cataract formation.

## INTRODUCTION

Connexins form intercellular channels, known as gap junctions, that enable direct cell-to-cell communication, allowing tissues to function as a syncytium by coordinating long-range electrical and metabolic activity^1,2^. These channels are critical in numerous physiological processes, and their dysfunction is linked to diverse diseases, including blindness, deafness, arrhythmia, stroke, neuropathy, skin disorders, and cancer^3–5^.

In addition to propagating cellular signals, gap junctions play a vital role in cellular stress responses. They dilute cytotoxic signals (e.g., potassium buffering by glial cells), and may also uncouple damaged cells from healthy tissue, preventing bystander effects or the so-called "kiss of death”^6,7^. Uncoupling is triggered by stress-induced gating mechanisms sensitive to transjunctional voltage and cytosolic chemical signals, like H^+^ and Ca^2+^ ^8–14^. Intracellular acidification, a hallmark of ischemia and cataract, is a universal regulator of gap junctional gating^15–18^. However, despite recent advances^19^, the molecular mechanisms underpinning pH-induced gating remain poorly defined.

Structurally, gap junctions are composed of twelve connexin subunits, forming either homomeric or heteromeric/heterotypic assemblies depending on isoform composition (21 isoforms in humans)^20,21^. Each hemichannel, formed by six connexins, docks with its counterpart in an adjacent cell to create a large-pore intercellular channel (∼1 kDa cutoff) capable of passaging ions, metabolites, and signaling molecules^22^. Each connexin subunit consists of four transmembrane helices (TM1–4), two extracellular loops (EC1–2) involved in junction formation, and an N-terminal domain (NT) implicated in channel selectivity and gating^23–26^ (see Fig. 1a). Intracellular loop (ICL) and C-terminal (CT) regions involved in trafficking and regulation are largely disordered and highly variable among isoforms^20,21^.

**Figure 1.**
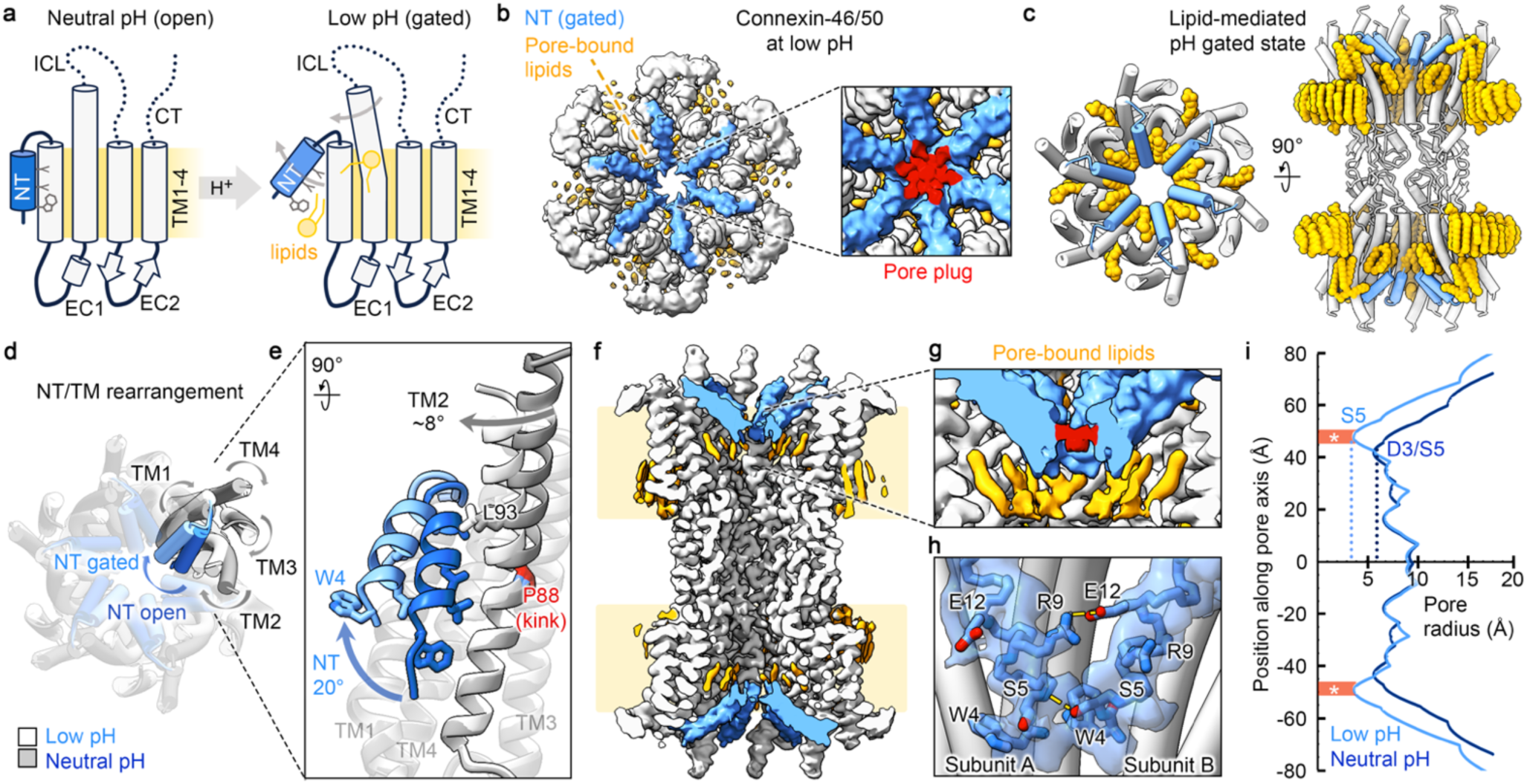
Low pH gated-state of connexin-46/50. **a** Schematic illustrating the pH-gating mechanism, where lipids stabilize the N-terminal domain (NT) in the gated-state upon acidification. **b** Cryo-EM map and **c** atomic model of Cx46/50 at low pH, highlighting the NT (light blue) and lipids (orange). Inset in panel b, shows a plug-like NT density observed at lower map thresholds (red), indicating complete pore occlusion. Lipids are observed within the pore lumen and at subunit interfaces, stabilizing the gated-state conformation. **d** Overlay of the previously described stabilized open-state (7JKC, TMs – grey, NT – dark blue) and the low pH gated-state (TMs – white, NT – light blue) showing concerted NT and TM movements induced by acidification**. e** Zoomed view of overlayed subunits, illustrating a 20° displacement of the NT toward the pore and rearrangement of hydrophobic residues (W4, L7 and L11). TM2 is displaced by approximately 8°, facilitated by a conserved proline kink (P88, red), maintaining hydrophobic contact between L93 (V93 in Cx50) and A14 (V14 in Cx50) on the NT. **f** Slice view of the cryo-EM map, showing lipids resolved within the channel pore, stabilizing the NT gated-state. **g** Inset, displaying the map at lower contour levels to highlight the pore plug and pore-bound lipid densities**. h** Close-up of NT gating domains with inter-subunit electrostatic interactions shown. The cryo-EM map for the NTs (blue transparency) showing fit of the model to the map features. **i** Pore profile plot comparing pore radii of low and neutral pH models, noting the proximal NT residues G2 and D3 are absent in the Cx46/50 low pH model. Regions associated with the unmodeled NT plug density are indicated (red; asterisk). Structural models display Cx46 as the representative isoform.

Early models of pH-gating based on studies of Cx43 and Cx50 proposed a "ball-and-chain" mechanism, linking the CT in pore occlusion^27–31^. However, subsequent studies have shown that pH gating in several isoforms, including Cx46 and Cx50, does not require the CT^32–35^. More recently, the NT domain has been implicated as a pH-gating particle, supported by cryo-electron microscopy (cryo-EM) studies of Cx26^19^. Though limited resolution of this study precluded atomic-level details of the gating mechanism, the study describes pH-induced conformational changes in the NT that result in pore occlusion. The NT’s involvement in pH-gating aligns with its conserved nature and its suggested role in other gating mechanisms, such as “fast” voltage-gating, underscoring its central function in connexin regulation^23,25,26,36–38^.

Recent cryo-EM studies of various connexin isoforms have revealed diverse NT conformations influenced by environmental conditions and suggest a potential role for lipids in modulating NT conformational states (recently reviewed^39^). However, the origination of pore-bound lipids in these studies remains unclear, as hydrophobic molecules can diffuse into the large-pore of these channels during detergent extraction or reconstitution into lipid nanodiscs, complicating interpretation of their potential functional relevance.

Here, using single-particle cryo-EM, we investigated the pH-gating mechanism of native Cx46/50 channels from aged lens tissue. We show that at neutral pH, channels predominantly adopt a stabilized open-state devoid of pore-bound lipids, consistent with prior studies^40,41^. When this sample is exchanged to mildly acidic conditions known to gate these channels in electrophysiology studies^32–35^, structural changes induce lipid infiltration into the pore, displacing the NT and driving an ensemble of gated states. Lipid infiltration is shown to originate from the surrounding lipid environment, to be essential for gating, fully reversible upon return to neutral pH, and is proposed to be linked to protonation of conserved histidines. Structural deconvolution reveals non-cooperative NT movements contributing to multiple gated states, alongside minor populations of stabilized and destabilized open-states. Collectively, these findings provide critical insights into the dynamic interplay between lipids and pH-gating in Cx46/50, offering molecular insights into age-related cataract formation and establishing a framework for understanding connexin regulation under cellular stress.

## RESULTS

### Low pH gated-state of connexin-46/50

Native Cx46/50 channels were purified from aged ovine lens tissue, containing native C-terminal truncations^42,43^, and reconstituted into MSP1E1 nanodiscs containing dimyristoyl-phosphatidylcholine (DMPC) lipids at pH 7.4, as previously described^41^. The reconstituted channels were then buffer-exchanged to pH 5.8 via gel filtration, to mimic low-pH conditions shown to gate C-terminal truncation variants of Cx46 and Cx50 channels in cellular functional studies^32–35^ (Extended Data Fig. 1,2). Cryo-EM single-particle analysis resolved a low-pH gated-state structure of Cx46/50 at 2.2 Å global resolution (Fig. 1b; Extended Data Fig. 3,4 and Table 1; Collection #1).

Heteromeric/-typic assembly patterns of Cx46/50 were indistinguishable from the cryo-EM density, consistent with prior studies^40,41^ and the >80% sequence identity across structured regions of the two isoforms. Atomic models for both isoforms were refined into a D6-symmetrized map, encompassing TM1-4, EC1-2, and the NT gating domain, while the CT and ICL regions remained largely unresolved due to intrinsic disorder (Fig. 1c; Extended Data Fig. 5 and Table 1). Over 400 lipids and 900 water molecules were also resolved, with extracellular leaflet lipids exhibiting long-range hexagonal packing, consistent with previous observations^41^. Given the similarity between Cx46 and Cx50, Cx46 is presented as the archetype, with isoform-specific differences noted where relevant (Extended Data Fig. 6).

The pH gated-state is characterized by inward displacement of the NT domain, as compared to the stabilized open-sate at neutral pH^41^, shifting 20° toward the pore center (Cα r.m.s.d. = 5.5 Å over the NT domains) (Fig. 1a,d,e; Extended Data Fig. 6). While the local resolution limited modeling of the NT to residue W4, a plug-like density at the center of the pore suggests full occlusion by proximal residues G2 and D3 (not explicitly modeled), consistent with complete channel closure (Fig. 1b inset,g,h). An additional constriction site at S5 narrows the pore in this region to 7.5 Å (compared to 11.6 Å in the open-state at D3), further illustrating the alteration of the NT in the pH-gated state (Fig. 1i).

The low pH gated-state of the NT is supported by infiltrated lipids, disrupting hydrophobic NT-TM1/2 interactions that anchor the NT in the stabilized open-state (Fig. 1a,c,g). The infiltrated lipids occupy a hydrophobic pocket formed by TM1/2 within the channel lumen, intercalated beneath the NT and stabilizing its displacement in the gated conformation. This gated-state appears further stabilized through a network of inter-subunit electrostatic interactions. Specifically, the S5 hydroxyl group forms a hydrogen bond with the backbone of F6 on a neighboring NT, while the E12 carboxylate forms a salt-bridge with R9 (N9 in Cx50) (Fig. 1h).

NT movement is coupled with a global clockwise rearrangement of the transmembrane helices, as viewed from the cytoplasmic side, that reinforce the gated NT conformation (Fig. 1d,e). TM2 exhibits the largest displacement, facilitated by a conserved proline kink at P88 that shifts the cytoplasmic portion of TM2 by

∼8° toward the pore, enabling hydrophobic interactions between L93 (V93 in Cx50) on TM2 and A14 (V14 in Cx50) on the NT (Fig. 1e). Additionally, subtle structural variation is observed compared to the open-state in the π-helix region of TM1 (residues ∼39–41) that forms a kink separating the para-helical region leading to EC1 (Extended Data Fig. 6).

### Lipid stabilization of the low pH gated-state

Beyond NT movement, the most striking difference between the open and gated-states is lipid translocation into the pore and at inter-subunit interfaces (Fig. 1c; Fig. 2). Two pore-bound lipids (PL1, PL2) form a double-layer hydrophobic “gasket” stabilizing the NT domain in the gated-state, while an interstitial lipid (IL) is identified intercalated at the subunit interface (Fig. 2a,b).

**Figure 2.**
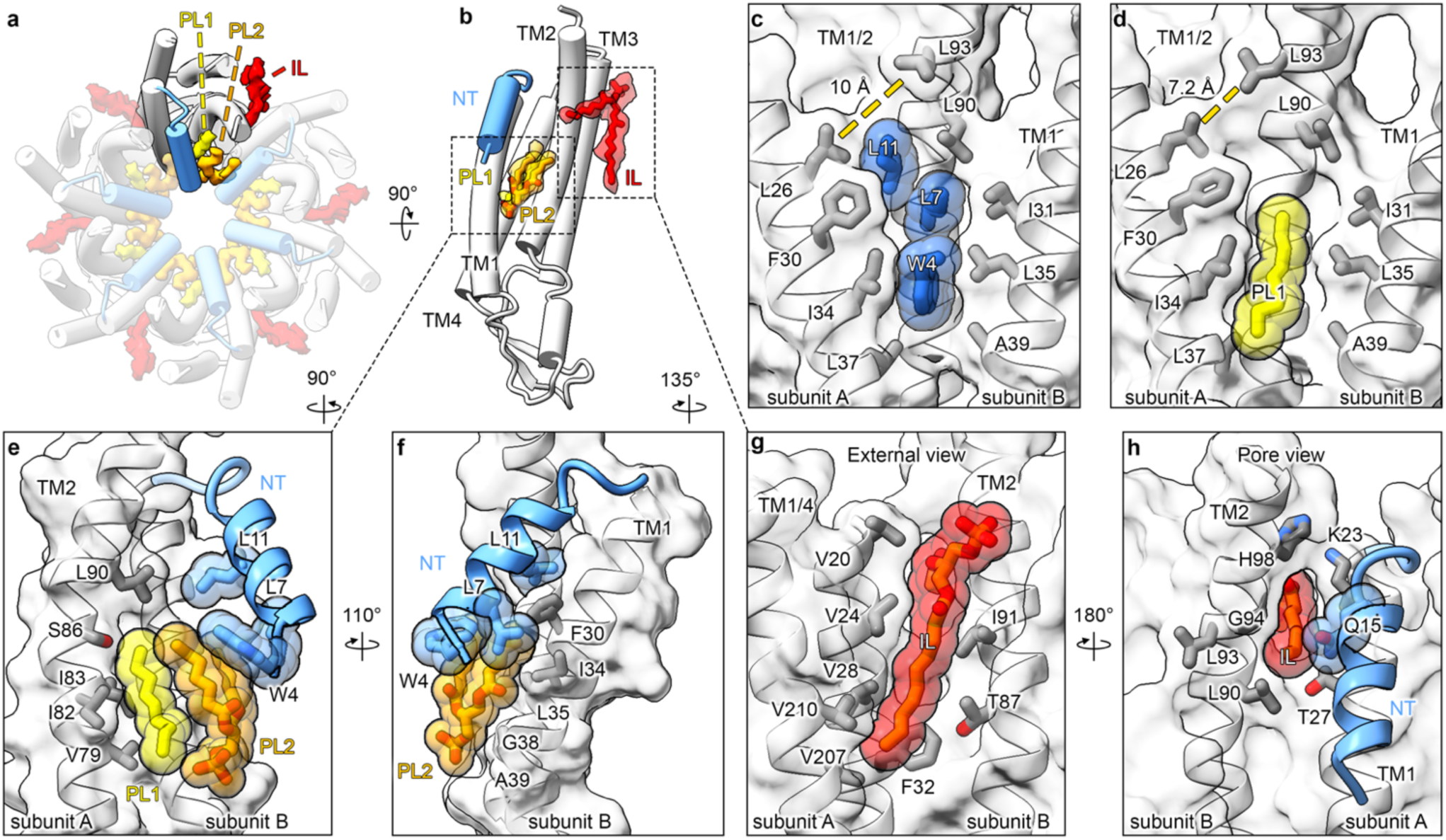
Lipid stabilization of the pH gated-state. **a** Cytoplasmic view of the Cx46/50 gated-state model showing segmented densities for the pore-bound lipids PL1 and PL2 (yellow and orange, respectively) forming a hydrophobic “gasket” that stabilizes the gated NT (light blue). Interstitial lipid (IL) densities (red) are intercalated between connexin subunits. **b** Single subunit view with modeled PL and IL lipids (stick representation) fitted to their respective segmented densities. **c** Zoomed view of Cx46 showing the NT hydrophobic residues (W4, L7 and L11) in the stabilized open-state at neutral pH (dark blue), anchored into the hydrophobic cleft formed by TM1/2 (grey sidechains). **d** At low pH, PL1 (yellow) occupies the NT anchoring site, and TM2 shifts 2.8 Å closer to TM1 compared to the neutral pH open-state (measured between L93 and L26, dotted lines). **e,f** Close-ups showing PL1 and PL2 inserted beneath the NT, supporting the gated-state through hydrophobic complementarity with TM1/2, with interacting amino acid sidechains labeled. **g,h** Views of IL intercalated between subunits, extending from the intracellular leaflet of the nanodisc lipid bilayer (g) toward the channel pore forming interactions with TM2 and the NT (h). Panels c-h display models of Cx46 as the representative isoform.

PL1, modeled as a nine-carbon acyl chain of DMPC, occupies a hydrophobic groove formed by TM1/2 of one subunit and TM1 of a neighboring subunit, displacing NT hydrophobic residues (W4, L7, and L11) that occupy this site in the open-state (Fig. 2c,d). The cytoplasmic portion of TM2 shifts toward the center of the pore under low-pH conditions, bringing L93 (V93 in Cx50) on TM2 ∼2.8 Å closer to L26 on TM1, contributing to NT displacement (Fig. 2c,d).

PL2 is positioned atop PL1 and forms additional interactions with TM1 of an adjacent subunit (Fig. 2a). Although the complete phosphocholine (PC) headgroup of PL2 could not be confidently modeled, the weak features extending from the U-shaped density suggest the headgroup extends toward the solvent, with the acyl chains oriented toward the hydrophobic pocket formed by PL1, TM1, and the NT (Fig. 2e,f). The hydrophobic anchoring residues of the NT rest on PL2, with the indole nitrogen of W4 positioned near the ester backbone of this lipid, potentially contributing to the stability of binding at this site (Fig. 2e,f).

The IL intercalated at the subunit interfaces aligns with the intercellular leaflet (Fig. 2a,b). The IL exhibits splayed acyl chains, with one chain forming extensive hydrophobic contacts with TM1/4 and TM2 of adjacent subunits and the other chain penetrating toward the NT and interacting with Q15 (N15 in Cx50) (Fig. 2g,h). These extensive interactions, particularly with TM2 and the NT, likely contribute to stabilizing the gated-state conformation. Notably, the IL density is absent in the open-state structure, suggesting a pH-dependent interaction and potential entry/exit site for lipid infiltration into the pore (discussed below).

### pH-gating is lipid dependent and reversible

We next sought to demonstrate that a lipid environment is required for pH-gating of Cx46/50 and that lipid recruitment into the pore is a reversible process driven by pH conditions. First, to determine lipid dependence, Cx46/50 channels were reconstituted into amphipol (PMAL-C8) at pH 7.4, following established methods for obtaining the open-state structure^40^, and then buffer-exchanged to pH 5.8. Cryo-EM single-particle analysis resulted in a 2.7 Å reconstruction, with NT domains resolved adopting the stabilized open-state conformation (Cα r.m.s.d. = 0.2 Å)^41^, with no lipids or detergent in the pore (Fig. 3a; Extended Data Fig. 2,6,7 and Table 1; Collection 2), confirming lipid dependence of the pH-gated state.

**Figure 3.**
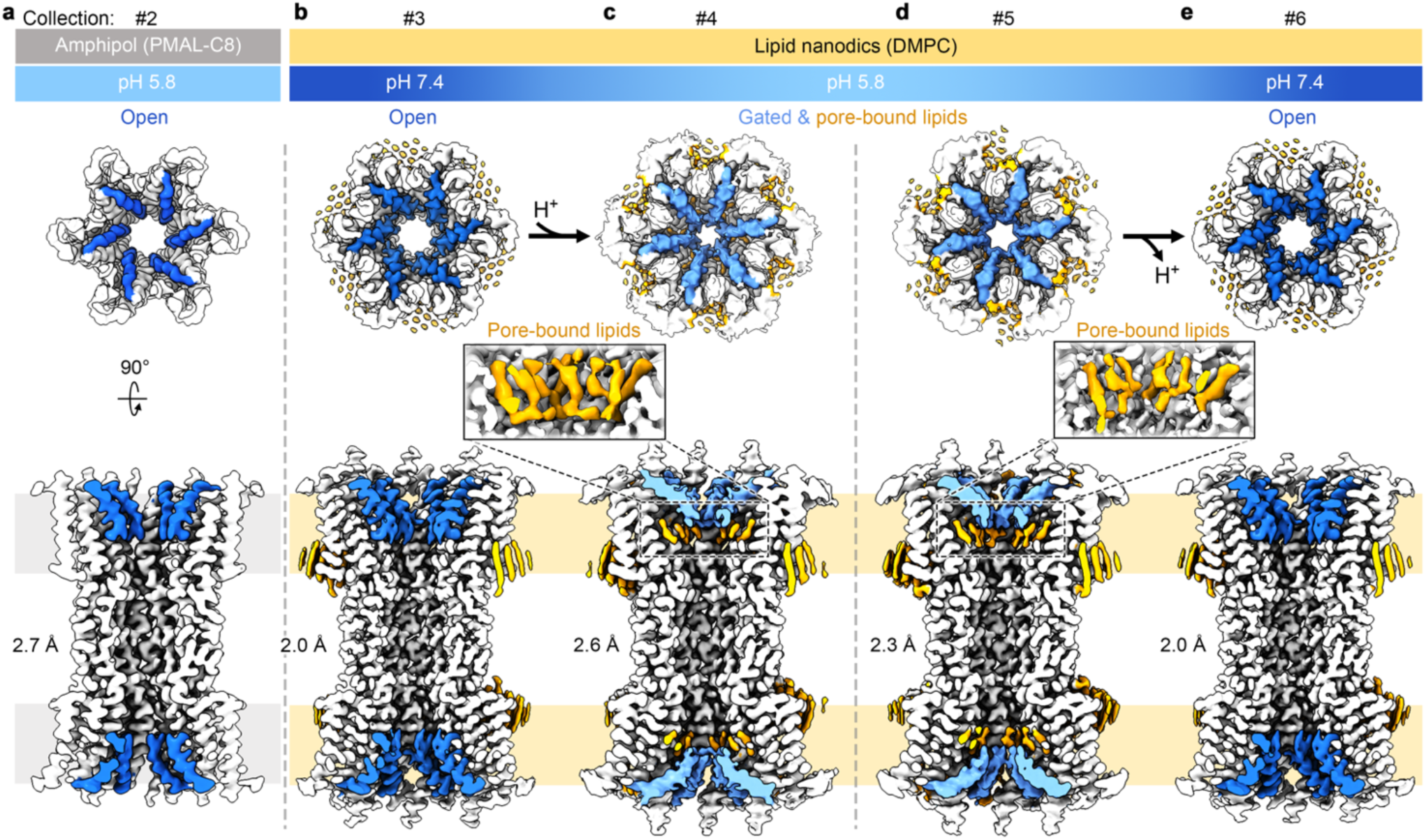
Lipid requirement and reversibility in the pH-gating mechanism. Cryo-EM maps of Cx46/50 channels under varying conditions, illustrating lipid-mediated and reversible pH gating. NT domains in open and gated-states are shown in dark blue and light blue, respectively, with lipids in orange. Specimens from different preparations are separated by vertical dashed lines. **a** Cx46/50 reconstituted in amphipol (PMAL-C8) at low pH, showing the NT in the open-state with no lipid (or detergent) densities observed in the pore. **b** Cx46/50 reconstituted into nanodiscs (containing DMPC lipids) and maintained in neutral pH conditions, showing the open-state with the pore free of lipids. **c** The same sample after exchange to low pH, resulting in the gated-stated with pore bound lipids (inset, highlights lipid densities). **d** Cx46/50 reconstituted into nanodiscs at neutral pH, then exchanged to pH 5.8 and maintained at low pH, showing the gated-state with pore-bound lipids (inset). **e** The same low pH sample returned to neutral pH resulting in the open-state and the pore is devoid of lipids.

Next, to ensure lipids enter the pore as a result of low pH-induced structural changes, rather than by nanodisc reconstitution artifacts, the following control experiments were conducted. Cx46/50 channels reconstituted into nanodiscs at pH 7.4 were divided into two samples and buffer-exchanged to either neutral (pH 7.4 control) or low pH (pH 5.8) conditions (Extended Data Fig. 2; Collections 3 and 4, respectively). Cryo-EM 3D reconstruction of the neutral pH sample yielded a 2.0 Å map with NT domains in the stabilized open-state and a pore free of lipids or detergent (Fig. 3b; Extended Data Fig. 8). In contrast, the low pH sample produced a 2.6 Å map, revealing PL lipids stabilizing the NT in the gated-state (Fig. 3c and *inset*), confirming that lipids are translocated from the local lipid environment in response to low pH-induced structural changes.

Finally, reversibility of lipid-mediated gating was tested by returning the low pH-gated channels to neutral pH conditions. Here, Cx46/50 channels reconstituted into nanodiscs at pH 7.4 were buffer-exchanged to pH 5.8 to induce the gated-state. This sample was then split into two groups: one remained at pH 5.8 during buffer exchange (pH 5.8 control), and the other was dialyzed back to pH 7.4 (reversed pH 7.4) (Extended Data Fig. 2; Collections #5 and #6, respectively). Cryo-EM reconstruction of the pH 5.8 control yielded a 2.3 Å map, confirming the NT gated-state with PL lipids (Fig. 3d and *inset*; Extended Data Fig. 9). In contrast, reconstruction of the reversed pH 7.4 sample produced a 2.0 Å map, showing the stabilized open-state conformation devoid of pore-bound lipids (Fig. 3e). These findings confirm that lipid translocation is reversible and essential for pH-dependent gating observed by cryo-EM.

### Putative role of conserved histidine pH-sensors

Having established pH gating is lipid-mediated, we next examined the potential mechanism underlying H⁺-induced conformational changes. Conserved histidine residues have been suggested to play a key role in the pH-dependent gating of several connexins^33,44–46^. These cytosolic sites correspond to H17 and H95 in Cx46/50 (Extended Data Fig. 6). The role of conserved histidine residues in pH gating are consistent with experimentally derived gating response pKa values of C-terminally truncated Cx46/50 (pKa ∼6.6 for Cx46^34^ and ∼7.1 for Cx50^32^) and the universal nature of pH sensing across connexin isoforms.

A mechanistic role for H95 in pH sensing had not been previously apparent, as it is positioned on TM2 facing the lipid environment. In the pH-gated state of Cx46/50, the imidazole ring of H95 lies within 3.0 Å of the negatively charged phosphate headgroup of the interstitial lipid (IL) (Fig. 4a). This proximity suggests that protonated H95 forms a salt-bridge with the IL, stabilizing the gated NT/TM2 conformation. Coulombic surface potential analysis shows a dramatic shift to positive potential at this binding site under acidic conditions (Fig. 4b,c). At neutral pH, deprotonation of H95 would disrupt this interaction (Fig. 4c), consistent with the absence of lipid density at this site in the open-state structures. These findings implicate H95 in recruiting the IL to its binding site and in stabilizing the gated state in a pH-dependent manner.

**Figure 4.**
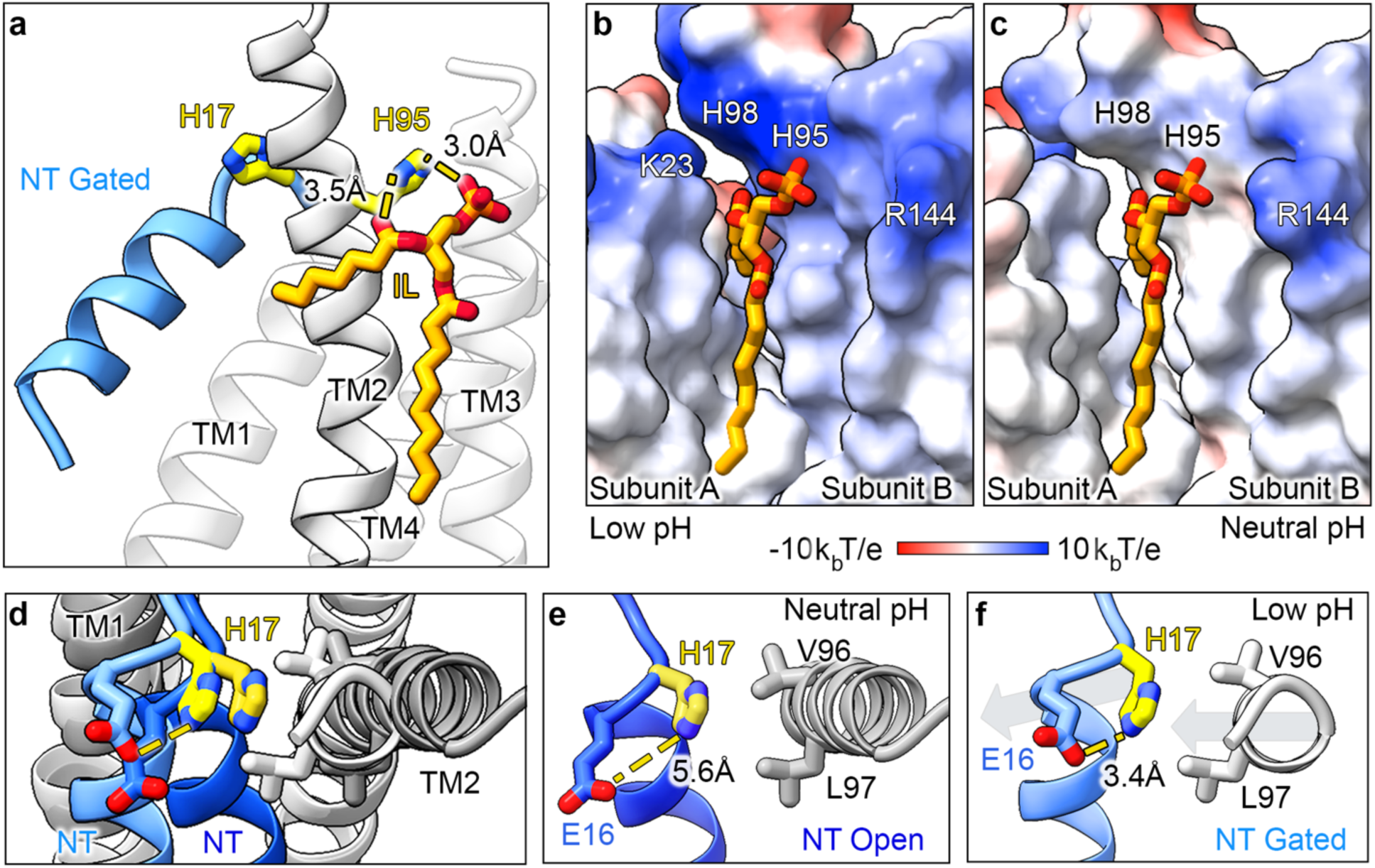
Putative roles of conserved histidine residues in pH-gating. **a** Zoomed view of a Cx46/50 subunit highlighting conserved histidine residues (yellow, stick representation): H17 on the NT (light blue) and H95 on TM2, oriented toward the lipid bilayer. The imidazole ring of H95 is within 3.0–3.5 Å of the negatively charged phosphate group and glycerol backbone of the interstitial lipid (IL; shown in orange, stick representation). **b** Zoomed view of the IL (orange, stick representation) intercalated at the Cx46/50 subunit interface (surface representation) colored by Coulombic potential (red – negative; white – neutral; blue - positive). The IL phosphate headgroup is accommodated by a positively charged cluster of positively charged amino acids, including the protonated states of H95 and H98, as well as K23 (R23 in Cx50) and R144 (R156 in Cx50). **c** The same IL binding site displayed with non-protonated states of H95 and H98. **d** Cytoplasmic view comparing the stabilized open-state (7JKC, NT – dark blue; TMs – gray) and gated-state (NT – light blue; TMs – white), showing correlated movements of the NT and TM2 involving rearrangements of E16 and H17. **e** At neutral pH, the imidazole ring of H17 is 5.6 Å from carboxylate sidechain of E16 and forms van der Waals interactions with V96 and L97 (A96 and V97 in Cx50) on TM2. **f** In the low pH gated-state, correlated movement of the NT and TM2 positions imidazole ring of H17 within salt-bridge distance (3.4 Å) of the negatively charged E16 sidechain. Panels b-f display models of Cx46 as the representative isoform.

Notably, mutating the equivalent H95 site does not eliminate pH sensitivity in Cx43^46^ or Cx46^33^, indicating additional pH-sensing residues contribute to gating. H17, another highly conserved histidine, has also been implicated as a pH-sensor in functional studies of Cx26^44^ and Cx36^45^. H17 resides at the NT-TM1 hinge near the channel entrance, at a similar z-position to H95 relative to the pore axis (Fig. 4a). At neutral pH, H17 interacts with V96 (A96 in Cx50) and L97 (V97 in Cx50) on TM2 (Fig. 4d,e). Under acidic conditions, these contacts are maintained, while correlated rearrangements of the NT and TM2 position the protonated sidechain of H17 within 3.4 Å of the negatively charged carboxylate group on E16, enabling formation of a salt-bridge (Fig. 4d,f). This stabilizing interaction may also contribute to reinforcing the NT gated conformation.

### Low pH induces an ensemble of gated- and open-states

Single-channel recordings show acidic conditions induce a complex “slow” gating transition, with channels fluctuating between closed and open states^33,47^. In contrast, at neutral pH and minimal transjunctional voltage, Cx46 and Cx50 channels predominantly adopt an open-state^26,31,48–50^. In agreement with this behavior, our structural analysis revealed pronounced conformational heterogeneity induced by low pH conditions, particularly within the NT.

To investigate the structural basis of this heterogeneity, and its potential relationship to the dynamic gating behavior, cryo-EM datasets for Cx46/50 at low pH were analyzed using 3D variability analysis (3DVA)^51^ (Fig. 5). The primary principal component revealed NT domains sampling both open and gated conformations symmetrically across the channel, accompanied by conformational changes in TM2 (Supplemental Movie 1), consistent with the structural rearrangements described above.

**Figure 5.**
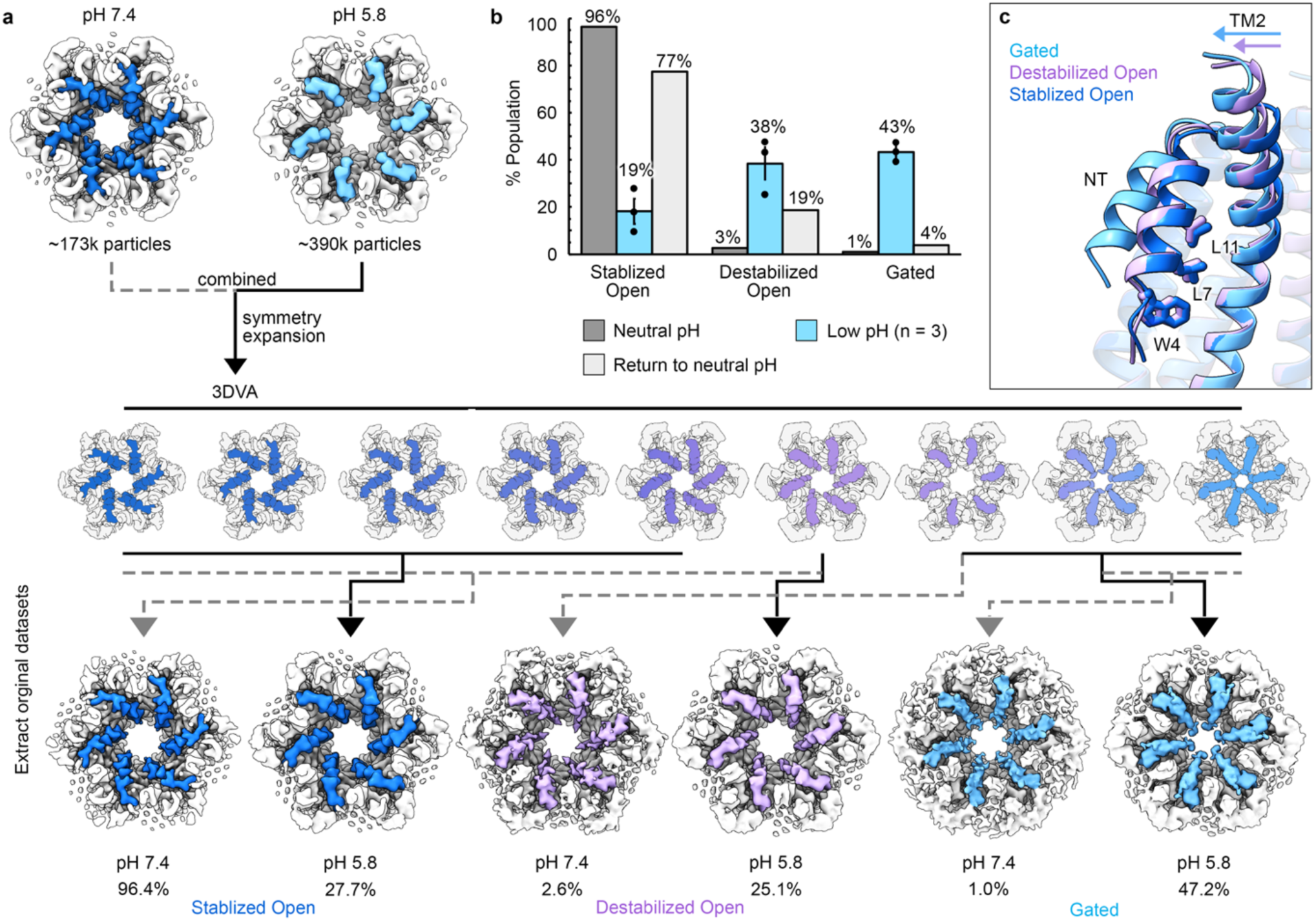
Low pH shifts the equilibrium ensemble of gated- and open-states. **a** Flow diagram summarizing the image processing strategy used to classify three distinct states of Cx46/50: stabilized open-state (NT – dark blue), destabilized open-state (NT – pink), and gated-state (NT – light blue), identified under neutral and low pH conditions. **b** Bar chart showing percent populations of the three states at neutral pH (gray, n = 1), after exchange to low pH (light blue, n = 3; mean ± s.e.m.), and after return to neutral pH following low pH exposure (light gray, n = 1). Low pH shifts the conformational equilibrium predominantly toward the gated-state. **c** Structural overlay of the three states, highlighting primary conformational differences localized to the NT and TM2 domains. The NT of the destabilized open-state shows slight displacement relative to the stabilized open-state, accompanied by a ∼6.4° movement of TM2 stemming from the P88 kink.

Particles corresponding to distinct states along the principal component were pooled for 3D reconstruction and used to estimate state distributions (Fig. 5a; see Methods). Across all low-pH datasets (Collections #1, #4, and #5), 43.2% ± 2.3% of particles were resolved in the lipid-mediated gated-state, while 18.2% ± 5.3% represented the open-state (mean ± s.e.m.; n = 3) (Fig. 5b). Notably, the open-state resolved at low pH was nearly identical to the stabilized open-state at pH 7.4 (Cα r.m.s.d. = 0.2 Å; Extended Data Fig. 6) and lacked pore-bound lipids.

In addition to the stabilized open and gated-states, a third state, termed the "destabilized open-state," was also identified along the same principal component, accounting for 38.5% ± 6.8% of low-pH particles (n = 3) (Fig. 5a,b). An atomic model built into a 3D reconstruction resolved to 2.2 Å (from Collection #1) showed the destabilized open-state resembles the stabilized open-state (Cα r.m.s.d. = 0.7 Å; Extended Data Fig. 6), but with subtle distinguishing differences in the NT and TM2 domains, as well as in the hydrogen bond patterns defining the TM1 π-helix kink at residues ∼39–41 (Fig. 5c; Extended Data Fig. 4-6).

In the destabilized open-state, the NT remains anchored to the lumen by hydrophobic residues (W4, L7, and L11) but adopts a distinct conformation, with a local Cα r.m.s.d. of 1.9 Å relative to the stabilized open-state NT. TM2 shifts ∼1.6 Å toward the pore center and bends by ∼6.4°, resembling but less pronounced than the movements of the gated-state (Fig. 5c). This conformation introduces a new pore-constriction site at S5, reducing the pore diameter to 9.7 Å in Cx46, which is narrower than the stabilized open-state (11.6 Å) but still sufficient for permeation of hydrated ions.

### Rare states detected at neutral pH

Previously, under neutral pH cryo-EM conditions, we have only detected the stabilized open-state of Cx46/50^40,41^. Consistent with this, 3DVA performed on neutral pH nanodisc datasets (Collections #3 and #6), as well as the amphipol low-pH dataset (Collection #2), revealed minimal NT heterogeneity, with variability limited to a slight “wobble” of TM2 (Supplemental Movie #). However, this approach may have lacked the sensitivity to detect rare conformational states within the dataset. Therefore, as an approach to enhance detection, we combined low-pH and neutral pH nanodisc datasets (Collections #3, #4 and #5, #6), ensuring a substantial representation of each state, and reanalyzed the combined particle set using 3DVA. This approach successfully resolved all three states (stabilized open, destabilized open, and gated) allowing for their classification and independent reconstruction from particles belonging only to the neutral pH dataset (Fig. 5a; see Methods).

This analysis further details the differences in state distributions between neutral and low pH datasets. At neutral pH, 87.0% ± 9.5% of particles were in the stabilized open-state, 10.7% ± 8.1% in the destabilized open-state, and only 2.4% ± 1.4% in the lipid-mediated gated-state (n = 2) (Fig. 5a,b). In the sample that was never exposed to low pH (Collection #3), the stabilized open-state accounted for 96.4% of particles, with the destabilized open-state and lipid-mediated gated-state representing 2.6% and 1.0%, respectively (Fig. 5a,b; Extended Data Fig. 8).

The distribution of open-state channels at neutral pH closely aligns with single-channel electrophysiology studies conducted under minimal transjunctional voltage^26,31,48–50^, underscoring the agreement between structural and functional analyses. In contrast, the observed proportion of low pH gated-state channels deviates from expectations from electrophysiology studies, which would predict a predominance of closed channels at the low pH conditions used for cryo-EM^32–35^. These discrepancies may stem from the absence of transjunctional voltage in cryo-EM conditions, as well as other methodological differences. Alternatively, they may indicate additional underlying heterogeneity in pH-gating transitions that was not fully captured by this approach (see below).

### Non-cooperative gating of the NT-domain

In addition to symmetric gating movements of the NT on both sides of the Cx46/50 intercellular channel, 3DVA of the low-pH datasets also revealed asymmetric gating movements of the NT domain, with each hemichannel operating independently (Collections #4 and #5) (Fig. 6a; Supplemental Movie 1). To further deconvolute the gated-state heterogeneity and to investigate potential cooperativity within hemichannels, cryo-EM density for individual subunits were isolated using symmetry expansion and signal subtraction, followed by 3D classification on the individual subunits. This approach distinguished subunits in the gated and open-states based on NT characteristics, though differences between stabilized and destabilized open-states could not be resolved (Fig. 6b; Extended Data Fig. 3,8,9). Among signal-subtracted subunits from low pH datasets, 67.0% ± 0.5% (mean ± s.e.m.; n = 3; Collections #1, #4 and #5) were identified in the gated-state. In contrast, the same workflow applied to neutral pH datasets (Collections #3 and #6) exclusively identified subunits in the open-state conformation (Extended Data Fig. 8,9).

**Figure 6.**
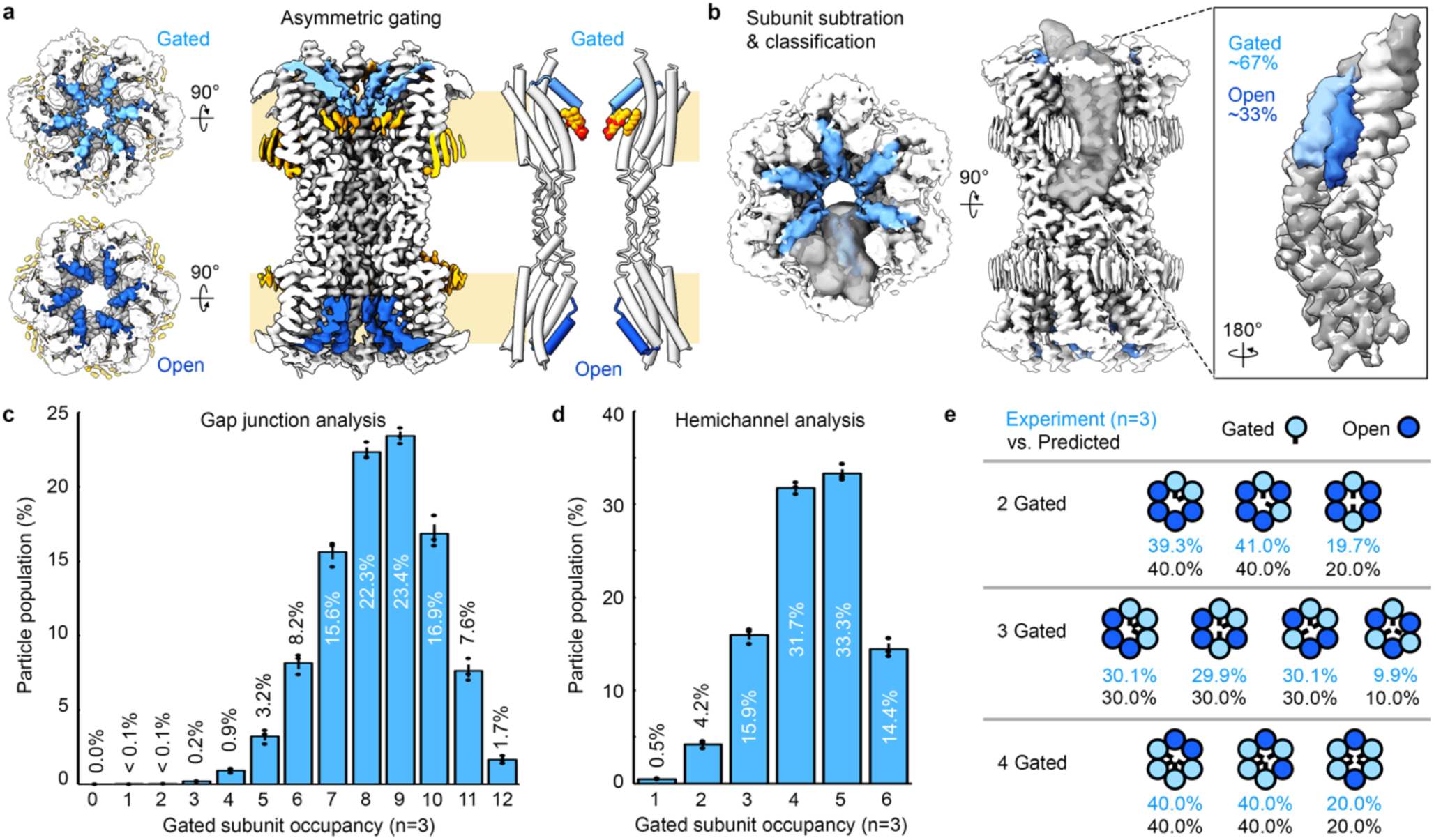
NT gating of Cx46/50 at low pH is non-cooperative. **a** Cryo-EM map and model of a an asymmetrically gated-state of the Cx46/50, where one side of the intercellular channel is gated (NT – light blue) with pore-bound lipids (orange), while the opposite side adopts the stabilized open-state. **b** Top and side views of the mask (grey) used for the symmetry expansion and signal-subtraction protocol, used to isolate density corresponding to the individual subunits for 3D classification. Inset, showing overlay of cryo-EM maps identified in the open-state (dark gray and dark blue NTs) and gated-state (light gray and light blue NTs) following signal-subtraction and 3D classification, where 67.0% ± 0.5% (mean ± s.e.m.; n = 3) were classified in the gated-state. **c** Bar chart showing the population of gated-state monomers identified at the gap junction level, and **d** hemichannel level. The populations display Boltzmann-like distributions, with the largest populations corresponding to nine gated subunits (23.4% ± 0.3%) at the gap junction level (panel c) and five gated subunits (33.3% ± 0.5%) at the hemichannel level (panel d). Bar charts show mean ± s.e.m. (n = 3), with individual data points displayed (black dots). **e** Possible configurations of open (dark blue) and gated-states (light blue) with quasi-equivalent number of gated subunits within each hemichannel. Each configuration is annotated with experimentally determined populations (mean; n = 3) compared to values assuming random distribution. The close correspondence of these values indicates NT gating at the subunit level is non-cooperative.

To quantify the frequency of open and gated-state monomers within parent gap junction particles, metadata from the symmetry expansion protocol were analyzed (see Methods). Remarkably, fully gated-states were rare, comprising only 1.7% ± 0.2% of gap junction particles and 14.4% ± 0.6% at the hemichannel level (mean ± s.e.m.; n = 3) (Fig. 6c,d). Subunit distributions in the gated-state followed a Boltzmann-like pattern, with nine gated subunits being the most frequent configuration (23.4% ± 0.3%; n = 3) (Fig. 6c). A similar trend was observed at the hemichannel level, with five gated subunits being most prevalent (33.3% ± 0.5%; n = 3) (Fig. 6d). States containing two to four gated subunits per hemichannel were observed in multiple configurations, and their populations closely matched predicted values for random distribution (Fig. 6e).

Notably, no fully open-state channels were observed under low-pH conditions, and < 0.5% were identified with 1–3 gated protomers. These findings suggest that NT pH-gating occurs independently between neighboring monomers under cryo-EM conditions, supporting a non-cooperative gating mechanism. Moreover, the prevalence of subunits adopting a gated conformation under acidic conditions more closely aligns with functional results, particularly considering models that suggest just a single subunit may be sufficient to initiate gating of gap junction channels^36^ (see Discussion).

### Comparison to other connexin structures

The pH-gated structure of Cx46/50 shares similarities as well notable differences with previously reported gap junction structures obtained under chemical gating conditions, such as Cx26 at pH 6.4^19^, Cx26 treated with CO_2_ maintained at pH 7.4^52^ and Cx43 at pH 8.0^53^. The TM and EC domains of Cx46/50 align closely with these structures, with Cα r.m.s.d. values (excluding the NT and cytoplasmic portion of TM2) ranging from 0.6–0.8 Å. In contrast, significant variations in NT conformations are evident across these structures; yet, the NT consistently contributes to the formation of the primary constriction site of the channel (Fig. 7a,b).

**Figure 7.**
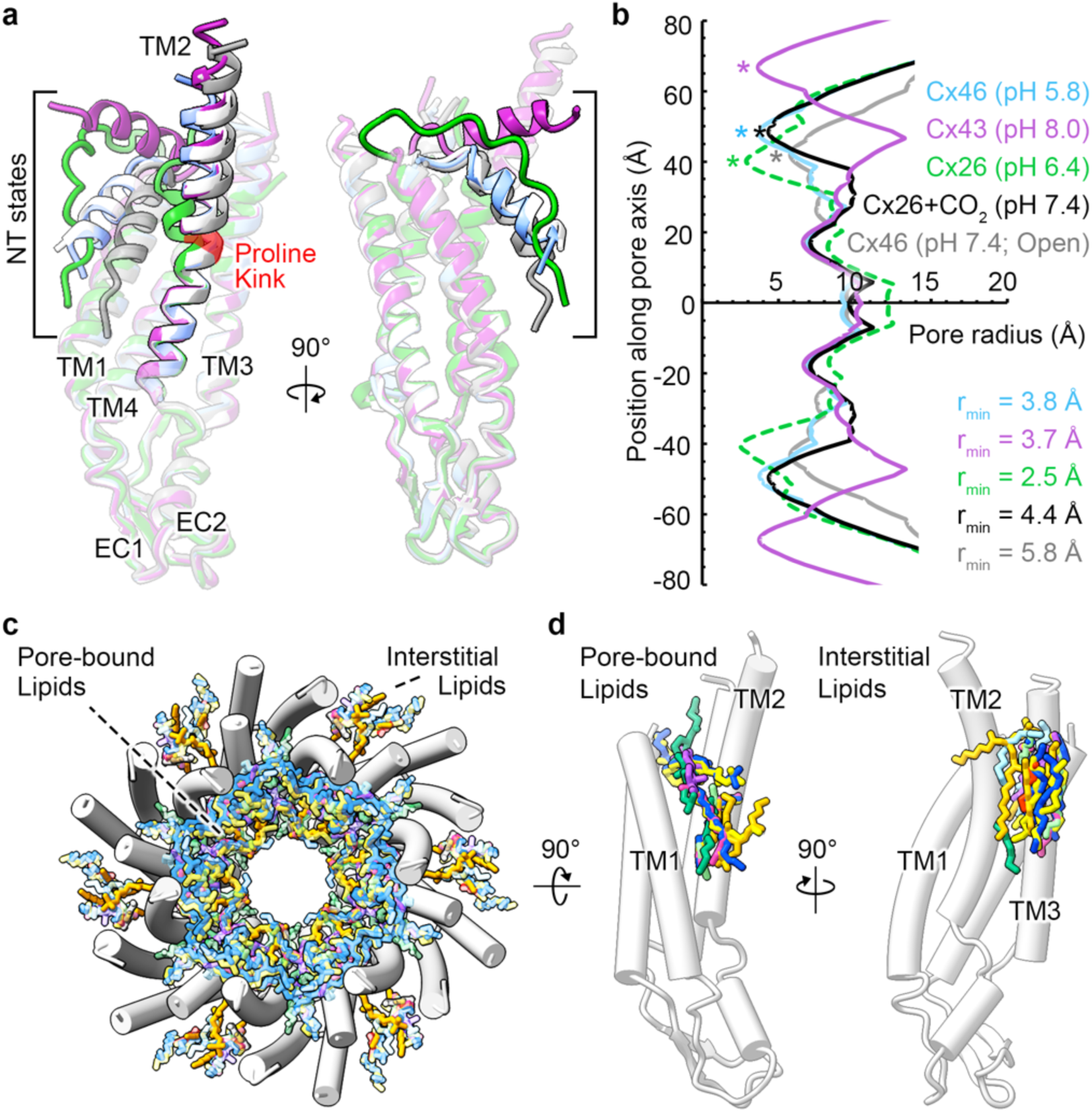
Structural comparison to connexin gap junctions under chemical-gating conditions. **a** Structural overlay of connexin subunits: Cx46 at pH 7.4 (7JKC, open-state, gray), Cx46 at pH 5.8 (gated-state, light blue), Cx26 at pH 6.4 (6UVT, green), Cx26 treated with CO_2_ maintained at pH 7.4 (7QEQ, white) and Cx43 at pH 8.0 (7F93, purple). The NT domain exhibits the most conformational variation, ranging from the stabilized open-state of Cx46 (pH 7.4), aligned along the pore axis, to the lifted NT conformation of Cx43, oriented orthogonal to the pore axis. The low pH gated-state NT of Cx46 (pH 5.8) most closely resembles the CO_2_–treated Cx26 structure (C*α* r.m.s.d. = 1.2 Å). TM2 shows differences stemming from a conserved proline kink (red). **b** Pore profile plot of the structures in panel a, colored accordingly. Asterisks indicate points of minimum radius (r_min_) for the modeled regions. The plot for Cx26 at low pH (6uvt, dotted line) reflects the absence of sidechains in the deposited model. **c** Top view of the Cx46 gap junction with the NT’s hidden, and superposition of lipids modeled in the channel pore of Cx46 at low pH (orange), Cx26 (7QEQ, red; 8QA0, grey), Cx36 (8IYG, green; 7XNH, light blue), and Cx43 (7F92, pink; 7XQF, yellow; 7XQB, blue; 7F93, light green), all sharing a similar trend of occupying portions of the channel lumen and at subunit interfaces. **d** *Left,* Cx46 subunits (NT hidden), showing superimposed pore-bound lipids (stick representation) from various connexin structures: Cx46 at low pH (orange), Cx36 (7XKT, green; 8IYG, purple), and Cx43 (7F92, pink; 7XQF yellow; 7XQB, blue; 7F93, green). *Right,* Cx46 subunit (NT hidden), showing superimposed annular lipids near the IL site from various connexin structures: Cx46 at low pH (orange), Cx26 treated with CO_2_ (7QEQ, red), Cx36 (7XKT, green; 8IYG, purple; 7XNH, light blue), and Cx43 (7F92, pink; 7XQF, yellow; 7XQB, blue; 7F93, green).

Among the compared structures, the NT of Cx26 treated with CO₂ at pH 7.4 is most similar to the pH-gated NT of Cx46/50 (Cα r.m.s.d. = 1.2 Å). In contrast, the largest deviation is observed in Cx43 (Cα r.m.s.d. = 12.2 Å), where the NT adopts a near-parallel orientation relative to the membrane bilayer. Interestingly, the NT of Cx46/50 shows substantial deviations with the pH-gated Cx26 structure (Cα r.m.s.d. = 9.1 Å), although their proximal positions indicate a shared role in pH gating (Fig. 7a,b). However, some discrepancy between these studies may reflect the limited resolution of the cryo-EM density used for modeling the Cx26 pH-gated state^19^.

Correlated with NT displacement, distinct conformational changes in TM2 were also observed. Compared to the stabilized open-state of Cx46/50, all other models display a more bent conformation localized to the cytoplasmic region of TM2 beyond the conserved proline kink (P88 in Cx46/50; P87 in Cx26 and Cx43) (Fig. 7a). When compared to this region of TM2 in the pH-gated state of Cx46/50, Cx43 at pH 8.0 showed the smallest deviation (Cα r.m.s.d. = 1.2 Å), followed by Cx26 treated with CO₂ (Cα r.m.s.d. = 2.2 Å), while Cx26 at low pH exhibited the largest deviation (Cα r.m.s.d. = 6.5 Å). These observations support a likely conserved mechanistic feature, where a hinge-like motion of TM2 is coupled to the conformational state of the NT.

Several recent connexin structures obtained under a variety of conditions have resolved lipids or detergents within the pore, though the source and conditions favoring their presence has been unclear (Fig. 7c,d)^39^. Similar to Cx46/50, lipids have been modeled at the NT docking site along TM1/2 in Cx36^54,55^ and Cx43^53^, where their presence correlates with NT displacement from the stabilized open-state observed in Cx46/50 (Fig. 7d). Additionally, stabilized lipids (or detergents) have been identified near the IL binding site of Cx46/50 in other connexin structures, including Cx26^52^, Cx36^54,55^, and Cx43^53^ (Fig. 7d), consistent with the relatively high conservation at these lipid binding sites (Extended Data Fig. 6). While the precise source of lipids from previous studies remains uncertain, these consistent observations suggest a conserved role for lipids in intercalating between subunits and modulating NT conformational states across diverse connexin isoforms.

## DISCUSSION

Using cryo-EM, we resolved a mosaic of native Cx46/50 gap junction structures under varying pH conditions, uncovering the structural basis of pH-gating (Fig. 8; Supplemental Movie 2). These findings reveal how NT domain conformations and lipid interactions may act synergistically to regulate intercellular communication. However, it is important to note that a direct role of lipids in gap junction gating remains to be validated through cellular functional studies. The observed insertion of lipids into the pore of other connexin isoforms, as well as related innexins^56^, pannexins^57^, and volume-regulated anion channels^58^, suggests this mechanism may have broad physiological relevance to large-pore ion channels. Thus, the dynamic role of lipids in stabilizing gating transitions in Cx46/50 may provide a broad mechanistic foundation for understanding tissue protection during ischemia, connexin-linked pathologies such as age-related cataract formation, and potential therapeutic strategies targeting connexin dysfunction in disease^59^.

**Figure 8.**
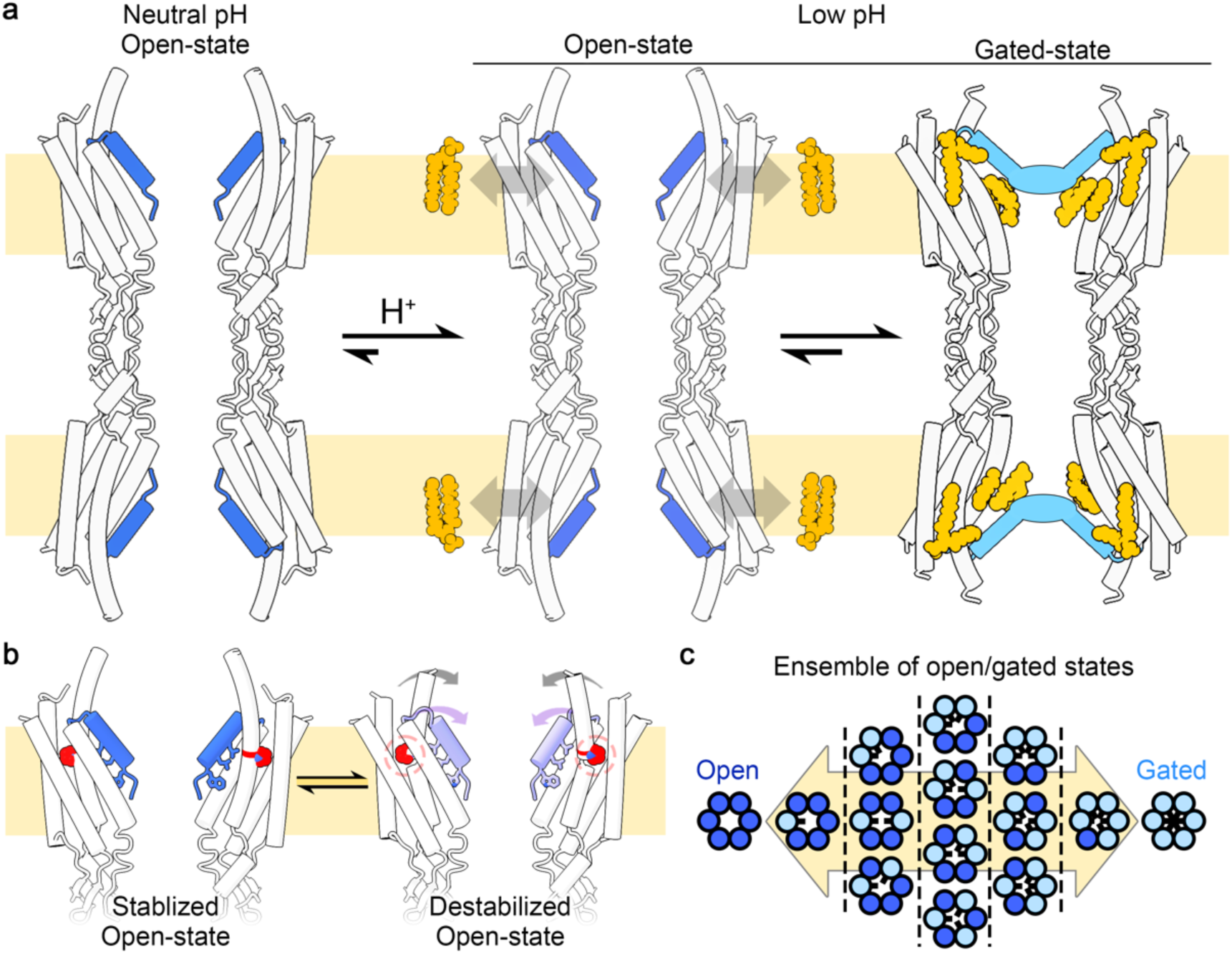
Schematic of reversible lipid-mediated pH gating of Cx46/50. **a** At neutral pH, Cx46/50 gap junctions predominantly adopt an open-state (NT – dark blue). At low pH, the equilibrium shifts toward an ensemble of open- and gated-states (NT – light blue), with the gated-state being most populated. Lipids (orange) translocate from the membrane environment into the pore, displacing the NT and occluding the channel permeation pathway. Upon returning to neutral pH, pore-bound lipids diffuse back to the membrane, and the NT returns to the open-state conformation. **b** The open-state represents a conformational equilibrium between a stabilized open-state and a less populated destabilized open-state (NT – pink), characterized by a slight displacement of the NT and bending of TM2 at a conserved proline kink (red). **c** The gated-state comprises a complex ensemble of micro-states, with NT domains of individual subunits adopting non-cooperative gated-state conformations.

### Lipid-mediated pH-gating mechanism of Cx46/50

At neutral pH, Cx46/50 channels predominantly adopt the stabilized open-state, devoid of pore-bound lipids or detergents. Through development of a deep-classification approach, we showed that this state is accompanied by minor populations of a destabilized open-state and a lipid-mediated gated-state at neutral pH (2.6% and 1.0% of particles, respectively) (Fig. 8a,b). Both open states maintain NT domain anchoring to TM1/2 via hydrophobic interactions, maintaining pore diameters sufficient for the permeation of hydrated ions.

Under mildly acidic conditions, the equilibrium shifts toward the lipid-mediated gated-state (Fig. 8a), with ∼67% of subunits adopting a gated conformation, while no fully open channels remain. Although the precise number of gated subunits required for pH-induced junctional uncoupling is unknown, functional studies suggest that a single subunit may be sufficient to trigger transjunctional voltage gating^36^. Thus, the observed cryo-EM ensemble may also align closely with electrophysiology studies which report a majority of channels become gated under similarly acidic conditions^32–35^. Notably, electrophysiology experiments are conducted in the presence of a transjunctional voltage potential, which may further drive closed-state occupancy.

Under cryo-EM conditions, low pH-induced gating is stabilized by pore-bound lipids (PLs) recruited from the annular lipid environment, filling hydrophobic sites that displace the NT, while interstitial lipids (ILs) at subunit interfaces reinforce TM2-NT interactions. In the absence of lipids, low pH had no structural effect on Cx46/50 (Fig. 3a), underscoring the essential role of lipids in gating. This lipid-mediated process is fully reversible, with lipids translocating back to the surrounding environment upon neutralization (Fig. 8a), aligning with the slow kinetics and reversibility observed in functional studies^33,34^.

Our findings, together with prior studies, highlight a coordinated movement of TM2 and the NT, facilitated by a conserved proline kink (P88) that functions as a hinge, conferring TM2 the flexibility to accommodate NT rearrangements in response to pH gating (Fig. 8a,b). The homologous proline in Cx26 and Cx32 has been previously linked to voltage-gating^60,61^, suggesting a broader conserved role for NT-TM2 coupling in mediating connexin gating across diverse physiological conditions.

### Asymmetric and non-cooperative gating

Our findings reveal that NT gating domains operate independently on either side of the intercellular channel, with asymmetrically gated particles resolved at low pH (Fig. 6a; Supplemental Movie 1). This state likely reflects physiological conditions in polarized gap junctions, where gating signals from one coupled cell selectively induces gating on one side of the channel^8,23^.

Further deconvolution of the structural ensemble revealed non-cooperative gating within individual hemichannels, with open and gated NT conformations distributed in a seemingly random pattern (Fig. 8c). This behavior aligns with functional studies on Cx32, which suggest that individual subunits can gate independently in response to transjunctional voltage^36^. Similarly, cryo-EM studies of Cx43 and Cx36 have identified non-cooperative NT conformations^53,54^. Collectively, these results support the notion that pH gating of Cx46/50 is more complex than a simple binary transition between fully open and fully closed states, involving an ensemble of gated-states. Future investigations will be necessary to define the minimal number of gated subunits required for complete junctional closure in response to pH.

### Protonation and lipid pathways

Conserved histidine residues, corresponding to H17 and H95 in Cx46/50, are shown to play key roles in pH sensing and gating in several connexins^33,44–46^. Our findings support these previous mutational studies, and suggest protonation of these conserved residues drive lipid interactions and stabilizes the NT in the gated-state, offering mechanistic insights into pH-dependent gating of Cx46/50 that may extend to other connexin isoforms. At low pH, H95 engages in a salt-bridge interaction with the phosphate headgroup of the IL, stabilizing the gated conformation through extensive hydrophobic interactions with the NT and TM2. Intriguingly, the position of the IL at the subunit interface, along with its pH-dependent recruitment to this binding site, suggest a potential pathway for entry or exit of PL lipids (Fig. 4a-c). Similarly, H17 undergoes a pH-dependent shift toward E16, forming a stabilizing salt-bridge with the NT in the gated state (Fig. 4d-f). While these interactions align with the proposed gating mechanism and underscore the importance of conserved histidine residues in pH-gating, further studies are needed to resolve the precise protonation events and explore the potential involvement of non-histidine residues, such as N-terminal acidic residues^62,63^. Additionally, as noted above, the physiological role of lipids in modulating connexin gating remains to be validated in cellular systems, including investigations into potential lipid specificity and translocation pathway into and out of the pore.

## ACKNOWLEDGEMENTS

We thank Dr. Thomas White for helpful discussions. We are grateful for instrumentation access and training provided by Dr. Claudia López and the staff at the OHSU Multiscale Microscopy Core, the OHSU Advanced Computing Center, and the Pacific Northwest Cryo-EM Center (supported by NIH Grant R24GM154185). The research was funded by NIH grant R35GM124779 (to S.L.R.)

## AUTHOR CONTRIBUTIONS

J.M.J. designed and executed the nanodisc experiments. J.B.M designed and executed the amphipol experiments. J.M.J. and J.B.M. collected the cryo-EM data. J.M.J. performed image processing and model building. J.M.J. and S.L.R. contributed to data analysis and prepared the initial manuscript. All authors contributed to final revision of the manuscript.

## METHODS

### Membrane scaffold protein expression and purification

A plasmid containing the sequence for a histidine tagged MSP1E1 was obtain from Addgene^64^ and the protein was expressed and purified as previously described with minor modification^41,65^. Freshly transformed *E.coli* cells (BL21 Gold-DE3, Agilent, Catalog #230130) were grown in LB medium supplemented with 50 μg mL^-1^ kanamycin at 37° C while being shaken at 250 r.p.m. After reaching an OD_600_ of 0.5-0.6, cultures were induced with 0.5 mM IPTG and allowed to express for 3 hrs post-induction at 37° C. Cells were harvested by centrifugation at 4000 x g for 20 min at 4° C and cell pellets were resuspended in ∼4 mL per gram of cell pellet of MSP Lysis Buffer (40 mM Tris pH 7.4, 1% Triton X-100, 1 mM phenylmethylsulfonyl fluoride (PMSF)). Cell suspensions were flash frozen in liquid nitrogen and stored at -80° C until needed.

Frozen cell suspensions were supplemented with 1 mM PMSF while thawing on ice and lysed by sonication. Crude lysate was cleared by ultracentrifugation at 150,000 x g for 20 min at 4° C. Sodium chloride was added to the supernatant for a final concentration of 300 mM. The supernatant was filtered (Millipore, 0.22 μm) and applied to a gravity column packed with HisPur Ni-NTA resin (Thermo Fisher Sci) that was prepared with Equilibration buffer (40 mM Tris pH 7.4 and 300 mM NaCl) at a ratio of 1 mL packed resin per gram of cell pellet. The supernatant was batch bound to the resin for 1 hr at 4° C with gentle rocking.

The MSP-bound resin was washed with 5 column volumes (CV) of Equilibration buffer, followed by 5 CVs of the following buffers: Triton buffer (40 mM Tris pH 8.0, 300 mM NaCl, 1% (vol vol^-1^) Triton X-100), Equilibration buffer, Cholate buffer (40 mM Tris pH 8.0, 300 mM NaCl, 50 mM cholate), Equilibration buffer, Imidazole Wash Buffer (40 mM Tris pH 8.0, 300 mM NaCl, 50 mM imidazole) and Equilibration buffer. MSP1E1 was eluted from the resin with 3 CV of Elution Buffer (40 mM Tris pH 8.0, 300 mM NaCl, 500 mM imidazole).

Eluate was concentrated to ∼5 mg mL^-1^ (UV_280_) using a 20 kDa m.w.c.o. spin concentrator (Vivaspin 6, Sartorius), filtered (Millipore, 0.22 μm) and applied to a gel filtration column (ENrich SEC 70; BioRad) using a fast protein liquid chromatography (FPLC) (NGC system; BioRad) equilibrated with 20 mM HEPES pH 7.4 and 150 mM NaCl. All chromatography steps were performed at 4° C. Peak fractions were monitored by UV_280_, pooled and concentrated to 300-500 μM using a 20 kDa m.w.c.o. spin concentrator. Samples were then aliquoted, flash frozen with liquid nitrogen and stored at -80° C until needed.

### Cx46/50 purification

Native Cx46/50 gap junctions were isolated from sheep lens fiber cells^40,41^. Fresh sheep eyes were obtained from Nebraska Scientific (Omaha, NE). Lenses were removed using a surgical blade and stored at -80° C. After thawing, the core lens fiber tissue, containing Cx46 and Cx50 intercellular channels that are C-terminally truncated (MP38)^42^ was dissected from the lens cortical tissue with a surgical blade. The lens core tissue was then Dounce homogenized and stripped membranes were prepared as previously described^40,41^. Stripped membranes were stored at -80° C in Storage Buffer (10 mM Tris pH 8.0, 2 mM EDTA, 2 mM EGTA) at a total protein concentration of 2 mg/mL (BCA, Pierce).

Thawed membranes were solubilized with 10 mM Tris pH 8.0, 2 mM EDTA, 2 mM EGTA, and 1% (wt vol^-1^) n-decyl-β-D-maltoside (DM) for 30 min at 37° C with gentle mixing. Insoluble material was cleared by ultracentrifugation 150,000 x g for 30 min at 4° C, followed by filtration (Millipore, 0.22 μm). The supernatant was then separated by anion exchange chromatography at 4° C (UnoQ, BioRad). Buffer A (10 mM Tris pH 8.0, 2 mM EDTA, 2 mM EGTA, 0.3% (wt vol^-1^) DM) was ran for 20 CV after sample application and protein was eluded with Buffer B (same a Buffer A with the addition of 1 M NaCl) using a gradient elution (to 35% Buffer B) over 20 CV. Peak fractions containing detergent solubilized Cx46/50 were concentrated to 150-300 μM as determined by UV_280,_ using a 50 kDa m.w.c.o. spin concentrator (Vivaspin 6, Sartorius).

### Reconstitution of Cx46/50 into lipid nanodiscs

Freshly prepared Cx46/50 was reconstituted at pH 7.4 into MSP1E1 nanodiscs using dimyristoyl-phosphatidylcholine (DMPC) lipids as previously described^41^, with minor modifications. Briefly, chloroform solubilized DMPC (Avanti) was dried under a stream of nitrogen gas creating a thin film and left under vacuum overnight to remove residual solvent. The DMPC thin film was solubilized in 20 mM HEPES pH 7.4, 150 mM NaCl and 5% (wt vol^-1^) DM to a final concentration of 35 mM. Detergent solubilized DMPC was then sonicated two times for 20 mins at 37° C with vortexing in between the sonication steps.

Solubilized Cx46/50 was mixed with the solubilized DMPC at a molar ratio of 0.6:90 (Cx46/50:DMPC) for 60 mins with gentle rotation at 37° C. Purified MSP1E1 was then added for a final molar ratio of 0.6:1:90 (Cx46/50:MSP1E1:DMPC) for 15 mins with gentle rotation at 37° C. Detergent was removed with an overnight incubation with SM-2 Bio-Beads (BioRad) at ratio of 30:1 beads:DM (wt:wt) with gentle inversion at 37° C. Bio-beads were removed by filtration and fresh beads were added for an additional 60 mins of gentle rotation at 37° C. The beads were removed by filtration and insoluble material was removed by centrifugation at 16,150 x g for 30 mins at 4° C.

The supernatant at pH 7.4 was filtered (Millipore, 0.22 μm) and applied to an FPLC gel filtration column (Superose 6 Increase 10/300 GL, GE Healthcare) equilibrated with Low pH buffer (20 mM Succinate pH 5.8, 150 mM NaCl, 2 mM EDTA and 2 mM EGTA). Peak fractions containing Cx46/50 embedded in nanodiscs were combined and concentrated using a 50 kDa m.w.c.o. spin concentrator (Vivaspin 6, Sartorius) to a final concentration of 1.8–2.2 mg mL^-1^ as determined by UV_280_. All chromatography steps were performed at 4° C.

### Reconstitution of Cx46/50 into amphipol

Freshly purified Cx46/50 was exchanged from DM to amphipol (PMAL-C8, Anatrace) as previously described^40^, with minor modification. Peak fractions containing Cx46/50 (determined by SDS-PAGE) from affinity purification (see above) were combined and applied to a gel filtration column (ENrich SEC 650, Bio-Rad) equilibrated with 20 mM HEPES pH 7.4, 150 mM NaCl, 2 mM EDTA, 2 mM EGTA and 0.3% DM (wt vol^-1^). Peak fractions containing DM solubilized Cx46/50 were pooled and combined with amphipol at a ratio 3:1 amphipol:Cx46/50 (wt:wt). The mixture was gently rotated overnight at 4° C, then Bio-beads were added at a ratio of 30:1 (wt:wt) beads:DM for 60 mins at 4° C with gentle rotation. The beads were removed by filtration and further cleared by centrifugation at 16,150 x g for 30 mins at 4° C. The supernatant containing Cx46/50 in amphipol at pH 7.4 was then filtered (Millipore, 0.22 μm) and then applied to a gel filtration column (ENrich SEC 650, Bio-Rad) equilibrated with Low pH buffer. Peak fractions containing Cx46/50 in amphipol were pooled and spin concentrated (Vivaspin, 50 kDa m.w.c.o., Satorius) to 1.3 mg mL^-1^. Protein concentration was determined by UV_280_ and sample purity was assessed by SDS-PAGE. All chromatography was done at 4° C.

### Controls for lipid mediated pH-gating

After nanodisc reconstitution of Cx46/50 and removal of Bio-beads at pH 7.4, as described above, the sample was then filtered (Millipore, 0.22 μm) and split in half. Half the reconstitution was applied to an FPLC gel filtration column (Superose 6 Increase 10/300 GL, GE Healthcare) equilibrated with Low pH buffer (20 mM Succinate pH 5.8, 150 mM NaCl, 2 mM EDTA and 2 mM EGTA). The other half was applied to a gel filtration column equilibrated with Neutral pH buffer (20 mM HEPES pH 7.4, 150 mM NaCl, 2 mM EDTA and 2 mM EGTA). For each pH condition, peak fractions containing Cx46/50 embedded in nanodiscs were combined and concentrated to a final concentration of 1.8-2.2 mg mL^-1^ as determined by UV_280_ using a 50 kDa m.w.c.o. spin concentrator (Vivaspin 6, Sartorius) and sample purity assessed by SDS-PAGE. All chromatography steps were performed at 4° C.

### Reversible lipid mediated pH-gating

To test whether lipid mediated gating was reversible, fresh Cx46/50 was reconstituted into nanodiscs at pH 7.4 as described above with the minor modification that after reconstitution the entire sample of Cx46/50 embedded in nanodiscs was applied to a gel filtration column (Superose 6, GE Healthcare) equilibrated in Low pH buffer (pH 5.8) to put the channels under gating conditions. Peak fractions containing Cx46/50 embedded in nanodiscs at low pH were pooled, divided in half, and then dialyzed (0.5–1 kDa m.w.c.o. Float-A-Lyzer G2; Spectrum Laboratories) overnight at 4° C with gentle stirring in either Neutral pH buffer or Low pH buffer (control). The ratio of dialysis buffer to sample was 100:1 (vol:vol). Dialysis buffer was exchanged after 2 hrs, and again after dialyzing overnight at 4° C with gentle stirring. After 2 hrs of dialysis with fresh buffer the following morning the sample was recovered and concentrated to 1.8–2.1 mg mL^-1^ as determined by UV_280_, using a spin concentrator (Vivaspin, 50 kDa m.w.c.o., Satorius).

### Negative stain electron microscopy

Cx46/50 embedded into lipid nanodiscs or solubilized with amphipol were prepared for negative stain EM, as previously described^40,41^. Briefly, 3 μL of sample (∼0.02 mg mL^-1^) was applied to a glow-discharged continuous carbon coated EM grid (Ted Pella, G400), blotted with filter paper and washed three times with gel filtration buffer. The sample was then stained with freshly prepared uranyl formate (0.75% wt vol^-1^, SPI-Chem). Negatively stained specimens were imaged on a 120 kV TEM (iCorr, Thermo Fisher Scientific) at a nominal 49,000 x magnification at the specimen level using a 2k x 2k CCD camera (Eagle 2 K TEM CCD, Thermo Fisher Scientific) with calibrated pixel size of 4.37 Å and with a nominal defocus of -1.5 μm, performed at OHSU’s Multiscale Microscopy Core.

### Cryo-EM specimen preparation and data collection

Samples were prepared for cryo-EM using 3 μL of freshly purified Cx46/50 embedded in nanodisc (1.8–2.2 mg mL^-1^) or amphipol (1.3 mg mL^-1^) applied to glow-discharged holey carbon grids (Quantifoil, R1.2/1.3 or R2/1). All cryo-samples were vitrified using a Vitrobot Mark IV (Thermo Fisher Scientific) at 20° C with 100% humidity and were blotted for 4–4.5 sec prior to being plunged into liquid ethane and stored in liquid nitrogen.

All cryo-EM specimens were imaged at the Pacific Northwest Cryo-EM Center (PNCC) on a Titan G4 Krios cryo-TEM (Thermo Fisher) operated at 300 kV. Movies for the amphipol dataset were recorded on a Gatan K2 summit direct electron detector at a nominal magnification of 29,000x (calibrated pixel size = 0.665 Å pixel^-1^) in counting mode, with an energy filter set to a 20 eV slit width. Nanodisc datasets were collected on a Gatan K3 direct electron detector at 29,000x or 105,000x nominal magnification (calibrated pixel size = 0.788 or 0.822 Å pixel^-1^) in super resolution mode, with energy filter slit widths of 10-20 eV. Data collections used 45 to 55 frames and total doses ranged from 40–50 e^-^ Å^-2^. Datasets of 555–12423 movies were collected using SerialEM^66^, with nominal defocus ranges of -0.8 to -2.2 μm or -0.5 to -1.8 μm. See Extended Data Table 1 for details of each dataset.

### Cryo-EM image processing

All datasets were processed in CryoSPARC (v4.4.1)^67^ and pre-processing was carried out in the same fashion across all datasets. Dose weighting and beam induced motion correction were done using Patch motion correction. Micrographs were binned to their nominal pixel size (2x super resolution pixel size) during Patch motion correction, here on referred to as the unbinned pixel size. Contrast transfer function (CTF) estimation was performed by Patch CTF estimation.

All datasets were processed using the same routine and the same masks used across datasets. Processing of Cx46/50 at low pH embedded in DMPC nanodiscs (Collection #1) will be described in detail as the representative dataset. Details for individual datasets are shown in the image processing workflow (Extend Data Fig. 3, 7-9). Using a subset of micrographs, semi-automated particle picking was used to create a preliminary 3D reconstruction for template particle picking on the entire dataset, resulting in 5,572,213 particle picks from 7,072 micrographs. After multiple rounds of 2D classification and removing duplicate particles, 685,022 bona-fide particles (binned to 2.4 Å pixel^-1^) were selected to generate an *ab-initio* model.

This was followed by a round of homogeneous refinement using D6 symmetry, unbinning of the particles (to 0.802 Å pixel^-1^) and a round of non-uniform refinement (D6 symmetry) leading to a 2.14 Å consensus reconstruction. The particles and map from the consensus refinement were used in heterogenous refinement with 6 classes (D6 symmetry) with a target resolution of 4 Å. Non-uniform refinements were then applied to all six classes, independently, using unbinned particle stacks and applying D6 symmetry. Particle stacks from 3D refinements that went below 3 Å (classes #0, #1 and #4, 476,719 particles total) were combined, down sampled (1.54 Å pixel^-1^, box 200 pixels) and subjected to another round of non-uniform refinement (D6 symmetry) resulting in a 3.16 Å resolution 3D reconstruction. The variability of this down sampled stack of particles was then assessed at the whole particle level by 3D Variability Analysis (3DVA)^51^ to guide the classification of various states for final 3D refinement.

### 3D variability analysis and classification of states

3DVA in CryoSPARC was used to parse through the conformational heterogeneity in the NT-gating domain at the gap junction level, and classify the gated-, destabilized open- and stabilized open-states. The particle stack from the binned non-uniform refinement (box 200 pixels, 1.54 Å/pix) was symmetry expanded using D6 symmetry yielding a particle stack of 5,720,628 particles prior to 3DVA. A custom 10 Å low-pass filtered mask that excluded the nanodisc was generated in CryoSPARC with an 8 Å resolution molmap created in ChimeraX using Cx46 (7JKC) that had additional residues placed into the pore to generate density to cover the channel entrances. 3DVA was performed using a target resolution of 4 Å, principal components (PC) of interest were selected and further analyzed. Ultimately in every dataset the primary PC was analyzed, as well as PCs that demonstrated asymmetric gating (Collections #4 and #5).

The intermediate 3DVA display job type was then used to classify the states of interest. The 3DVA display job was used to separate the PC into nine frames by setting the rolling window to zero and outputting particle subsets, generating unique stack of particles along the PC of interest. Each frame’s particle stack was independently locally refined (D6 symmetry) using the same mask used for 3DVA, duplicate particles were removed, and then particles were unbinned for a final round of non-uniform refinement with D6 symmetry applied. Particle stacks yielding reconstructions of similar states (i.e., similar NT conformation) were then combined. Three distinct states were identified (gated, destabilized open and stabilized open) and used for model building. The stabilized open-state map had a final resolution of 2.5 Å with 42,646 particles, the destabilized open-state map had a final resolution of 2.2 Å with 218,088 particles, and the gated-state map had a final resolution of 2.2 Å with 198,875 particles (gold-standard FSC) (Extended Data Table 1).

### Merging cryo-EM datasets for identification of rare neutral pH states

To computationally isolate rare states at neutral pH, datasets from neutral and low pH conditions were merged and subjected to 3DVA. This was achieved by merging cryoEM datasets samples that were prepped from the same specimen, collected on the same cryo-electron microscope with the same settings (*i.e.,* Collections #3 and #4 were merged and Collections #5 and #6 were merged). As an example, Collections #5 and #6, belonging to the reversible lipid mediated pH-gating experiment will be described in detail. The unbinned particle stacks (0.8016 Å pixel^-1^, box 384 pixels) obtained after heterogenous refinement from the respective datasets were merged. The low pH control dataset (Collection #5) contributed 764,205 particles and the neutral pH particle stack (Collection #6) contributed 427,873 particles, for a total of 1,192,078 particles.

The merged unbinned particle stack was down sampled (box 160, 1.97 Å/pix) and subjected to a round of non-uniform refinement (D6 symmetry), and then symmetry expanded using D6 symmetry for a total of 14,304,936 particles prior to 3DVA. The symmetry expanded particles were then analyzed by 3DVA with a target resolution of 4 Å, using the same mask used for the 3DVA on the independent datasets described above. PCs of interest were split into 9 frames/particle stacks along a PC using the intermediates 3DVA display output. Each particle stack was independently locally refined using particles from both low and neutral pH datasets. The particles were then reverted to their original datasets, and then deduplicated to undo symmetry expansion.

The particles were then unbinned and individual particles stacks were subjected to non-uniform refinement (D6 symmetry). Particles stacks demonstrating similar states were pooled and a final round of non-uniform refinement (D6 symmetry) was performed. Using this approach, all three states (gated, destabilized open and stabilized open) were identified in the neutral pH datasets revealing rare populations of gated-state gap junctions at neutral pH (1% for Collection #2 and 4% for Collection #5), which were previously not observed in these datasets using other common approaches of 3D classification. To assess for influence of initial map bias on end state maps, initial maps of open and gated states and initial low pass filters (16 Å) were both assayed as starting models, yielding similar results.

### Subunit classification

Subunit-level heterogeneity was analyzed using signal subtraction and 3D classification in CryoSPARC. This workflow was applied to particles from Collections #1, #3, #4, #5 and #6. The description here focuses on the workflow for Collection #4, as the exemplar approach.

A non-uniform refinement particle stack containing 476,719 high-resolution particles was down sampled by 2x binning (1.6 Å pixel^-1^, 240-pixel box size). A subset of 100,000 randomized particles (∼21% of the total) was symmetry expanded (D6 symmetry), generating 1.2 million particles that were then subjected to local refinement using D6 symmetry. A mask for signal subtraction was generated in ChimeraX by creating an 8 Å molecular map (molmap), generated from overlaid monomer models of the stabilized open and gated states. After signal subtraction, particles were re-centered and subjected to 3D classification without alignment. Classification parameters were optimized as follows: three classes, a target resolution of 3 Å, an O-EM learning rate of 0.3, an O-EM batch size of 2,000 particles, no per-particle scale factors, an initial low-pass filter resolution of 8 Å, and hard classification. Open and gated states were distinguished based on the position of the NT gating domain.

To determine the distribution of open and gated-state monomers within gap junction particles, metadata from the signal-subtracted 3D classified particles was analyzed. The following metadata fields were extracted into a CSV file from each class’s exported particle .cs file: uid, sym_expand/idx, and sym_expand/src_uid. Metadata from similar classes were combined to generate concatenated files, representing open and gated states, respectively.

The frequency of open and gated-states within gap junction particles and hemichannels was analyzed independently. An in-house developed bash script, “*Subunit State Frequency*”, was used to calculate the frequency of ‘sym_expand/src_uid’ occurrences to determine the distribution of open and gated monomers in gap junctions. For example, if a gated monomer’s ‘sym_expand/src_uid’ appeared 4 times, then 4 of the 12 subunits in the gap junction were classified as gated. To analyze hemichannels, another custom bash script, “*Gap Junction Splitter*”, was employed to split gap junction particles into their respective hemichannels based on the ‘sym_expand/idx’ metadata. The resulting hemichannel metadata was then processed using the same *Subunit State Frequency* script to evaluate the distribution of open and gated monomers within the individual hemichannels of the parent gap junction particles. Frequency data for both gap junctions and hemichannels were subsequently plotted to visualize the distribution of gated states across the dataset.

To investigate potential cooperativity of subunit gating across hemichannels within gap junction particles, a custom bash script, “*Hemichannel Arrangement*”, was developed to identify the relative positions of open/gated conformers and applied to the gated-state particle metadata. Initially, gap junction particle metadata was split into hemichannels using the *Gap Junction Splitter* script described above. Subsequently, the metadata for each hemichannel was analyzed independently to evaluate the relative positions of gated subunits in hemichannels containing two, three, or four gated subunits. The *Hemichannel Arrangement* script determined the number of gated subunits in each hemichannel by counting the occurrences of unique ‘sym_expand/src_uid’ IDs. For hemichannels with two to four gated subunits, the script analyzed the specific arrangements of gated subunits based on the combination of ‘sym_expand/idx’ IDs. Each unique arrangement was tallied to quantify the distribution of gated-state configurations within the hemichannel metadata. These results provided insights into the spatial organization and potential cooperativity of subunit gating across the hemichannels.

### Atomic modeling and validation

Previously published models of Cx46 (7JKC) and Cx50 (7JJP) in the stabilized open-state were used as starting points for atomic modeling in Coot^68^. For the gated and destabilized open-states, the NT domains were pruned to the 20^th^ residue and rebuilt into the respective maps. To avoid bias, all histidine were modeled in the protonated state for low pH datasets. Models were then iteratively refined in Phenix^69^ and Coot until convergence of statistics determined by MolProbity^70^. After the Cx46 models statistics converged, point mutations were introduced in Coot to convert to the Cx50 isoform and the model refinement/validation process was repeated until convergence (Extended Data Table 1).

Structural analysis and visualization of Coulombic surface potentials were performed in ChimeraX^71^. Pore profiles were analyzed using the program HOLE^72^. For this analysis, the sidechain of R9 on Cx46 (7JKC) was pruned to C*β* to account for the dynamic nature of this residue as demonstrated *in silco*^26^ and the weak density for the sidechain in the original cryo-EM map^41^. Sequence alignments were created using Clustal-w^73^ and visualized in Jalview^74^.

### AI-assisted technologies

During the development of this work ChatGPT (OpenAI) was used for help in writing code and to revise portions of the text to improve clarity. After using this tool for coding, synthetic datasets were generated and used for testing and validation. The authors reviewed and edited any content that was generated with this tool and take full responsibility of the content of this publication.

## DATA AVAILABILITY

Cryo-EM maps have been deposited to the Electron Microscopy Data Bank under accession codes EMDB-XXXXX (stable-open Cx46/50 at pH 5.8), EMDB-XXXXX (destabilized-open Cx46/50 at pH 5.8), EMDB-XXXXX (gated Cx46/50 at pH 5.8), EMDB-XXXXX (stable-open Cx46/50 at pH 7.4), EMDB-XXXXX (destabilized-open Cx46/50 at pH 7.4), EMDB-XXXXX (gated Cx46/50 at pH 7.4), EMDB-XXXXX (asymmetric gated Cx46/50 at pH 5.8) and EMDB-XXXXX (open Cx46/50 at pH 5.8 in amphipol). Coordinates for Cx46 atomic models have been deposited to the Protein Data Bank under accession codes XXXX (stable-open Cx46 at pH 5.8), XXXX (destabilized-open Cx46 at pH 5.8), XXXX (gated Cx46 at pH 5.8), XXXX (stable-open Cx46 at pH 7.4), XXXX (destabilized-open Cx46 at pH 7.4), XXXX (gated Cx46 at pH 7.4), XXXX (asymmetric gated Cx46 at pH 5.8) and XXXX (stable-open Cx46 at pH 5.8 in amphipol). Coordinates for Cx50 atomic models have been deposited to the Protein Data Bank under accession codes XXXX (stable-open Cx50 at pH 5.8), XXXX (destabilized-open Cx50 at pH 5.8), XXXX (gated Cx50 at pH 5.8), XXXX (stable-open Cx50 at pH 7.4), XXXX (destabilized-open Cx50 at pH 7.4), XXXX (gated Cx50 at pH 7.4), XXXX (asymmetric gated Cx50 at pH 5.8) and XXXX (stable-open Cx50 at pH 5.8 in amphipol). The original multi-frame micrographs have been deposited to EMPIAR under accession codes EMPIAR -XXXXXX (Collection #1, Cx46/50 at pH 5.8 in nanodiscs), EMPIAR - XXXXXX (Collection #2, Cx46/50 at pH 5.8 in amphipol), EMPIAR -XXXXXX (Collection #3, Cx46/50 at pH 7.4 in nanodiscs), EMPIAR -XXXXXX (Collection #4, Cx46/50 at pH 5.8 in nanodiscs), EMPIAR - XXXXXX (Collection #5, Cx46/50 at pH 5.8 in nanodiscs) and EMPIAR -XXXXXX (Collection #6, reversibly gated Cx46/50 at pH 7.4 in nanodiscs).

## CODE AVAILABILITY

In-house code developed for the analysis of image processing meta data have been deposited on the Reichow Lab GitHub repository (https://github.com/reichow-lab/subunit_analysis).

## EXTENDED DATA (TABLES, FIGURES AND LEGENDS)

**Extended Table 1.**
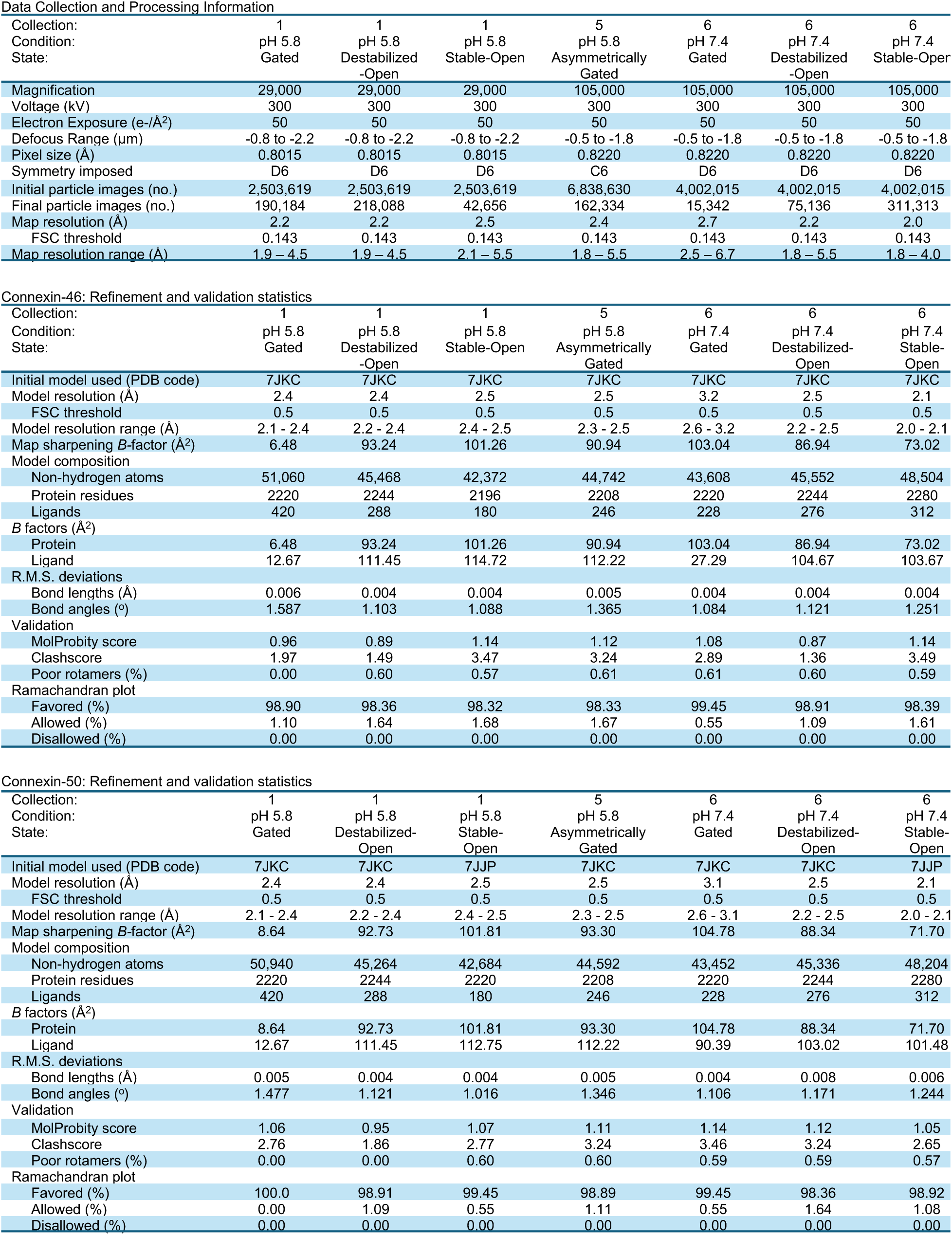
Cryo-EM data collection, refinement, and validation statistics.

**Extended Data Figure 1.**
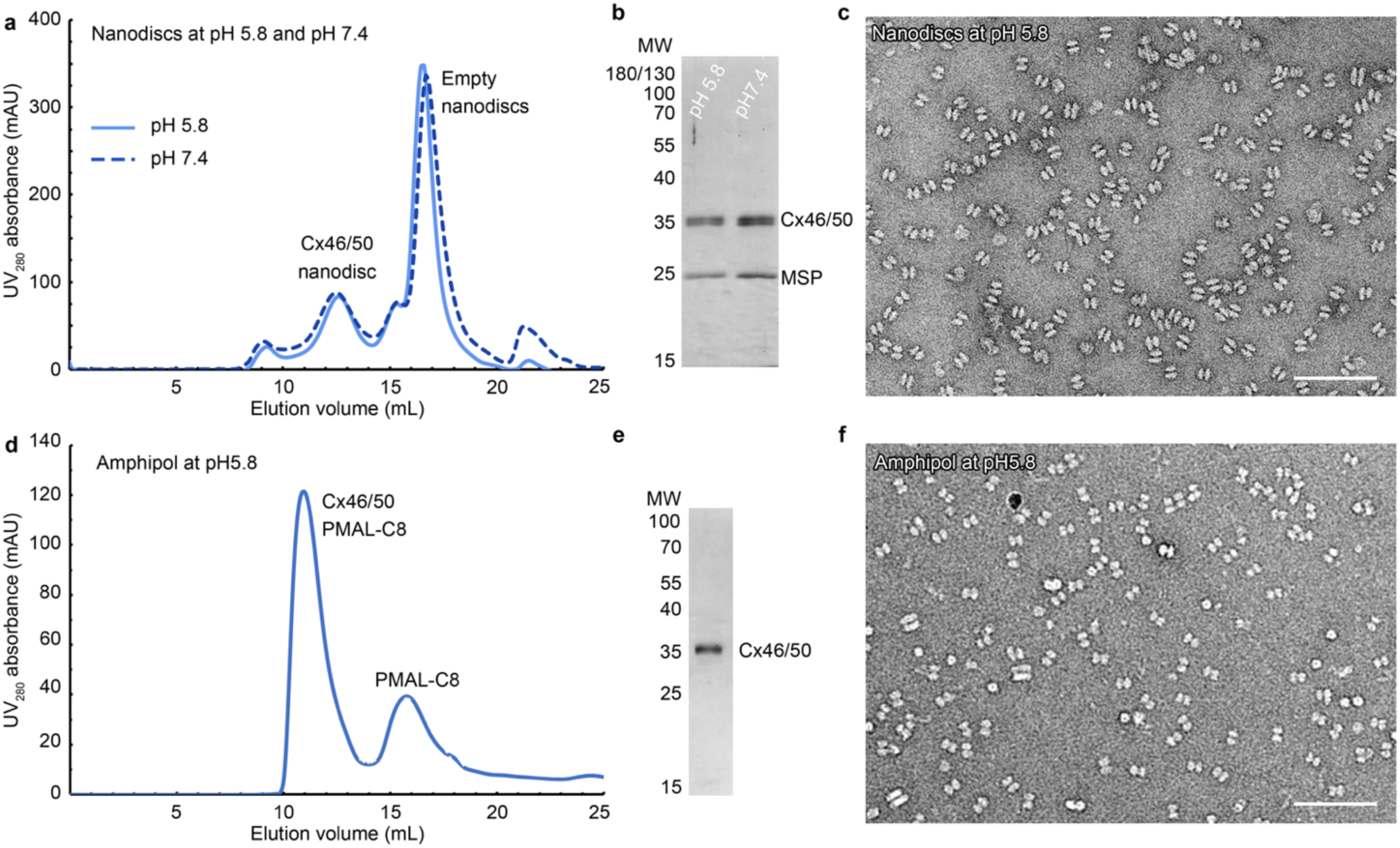
Purification and reconstitution of connexin-46/50 in nanodiscs and amphipols. **a** Gel filtration chromatography (Superose S6) traces monitored by UV absorbance at 280 nm. Peaks containing native Cx46/50 gap junctions embedded in DMPC lipid nanodiscs (MSP1E1) and empty nanodiscs are labeled. Gel filtration traces at pH 5.8 (light blue) and pH 7.4 (dashed, dark blue) are nearly identical. **b** Silver stained SDS-PAGE of the gel filtration peak containing Cx46/50 embedded in nanodiscs, with molecular markers indicated. MSP1E1 migrates at ∼24 kDa (predicted MW ∼27.5 kDa), and native Cx46/50 gap junctions from the lens core migrate together as a band at ∼38 kDa. **c** Electron micrograph of negatively stained particles from the pH 5.8 gel filtration fraction, containing Cx46/50 in nanodiscs (scale bar = 100 nm). Structures characteristic of Cx46/50 in nanodiscs are observed^41^. **d** Gel filtration chromatography (Bio-Rad ENrich SEC 650) trace monitored by UV absorbance at 280 nm. Peaks corresponding to native Cx46/50 reconstituted into amphipol (PMAL-C8) and PMAL-C8 micelles are labeled. **e** Silver stained SDS-PAGE of the gel filtration peak containing Cx46/50 in PMAL-C8 (band at ∼38kDa), with molecular weight markers indicated. **f** Electron micrograph of negatively stained particles from the pH 5.8 gel filtration fraction, showing Cx46/50 in amphipol (scale bar = 100 nm). Particles appear identical to those previously reported for Cx46/50 in amphipol at pH 7.4^40^.

**Extended Data Figure 2.**
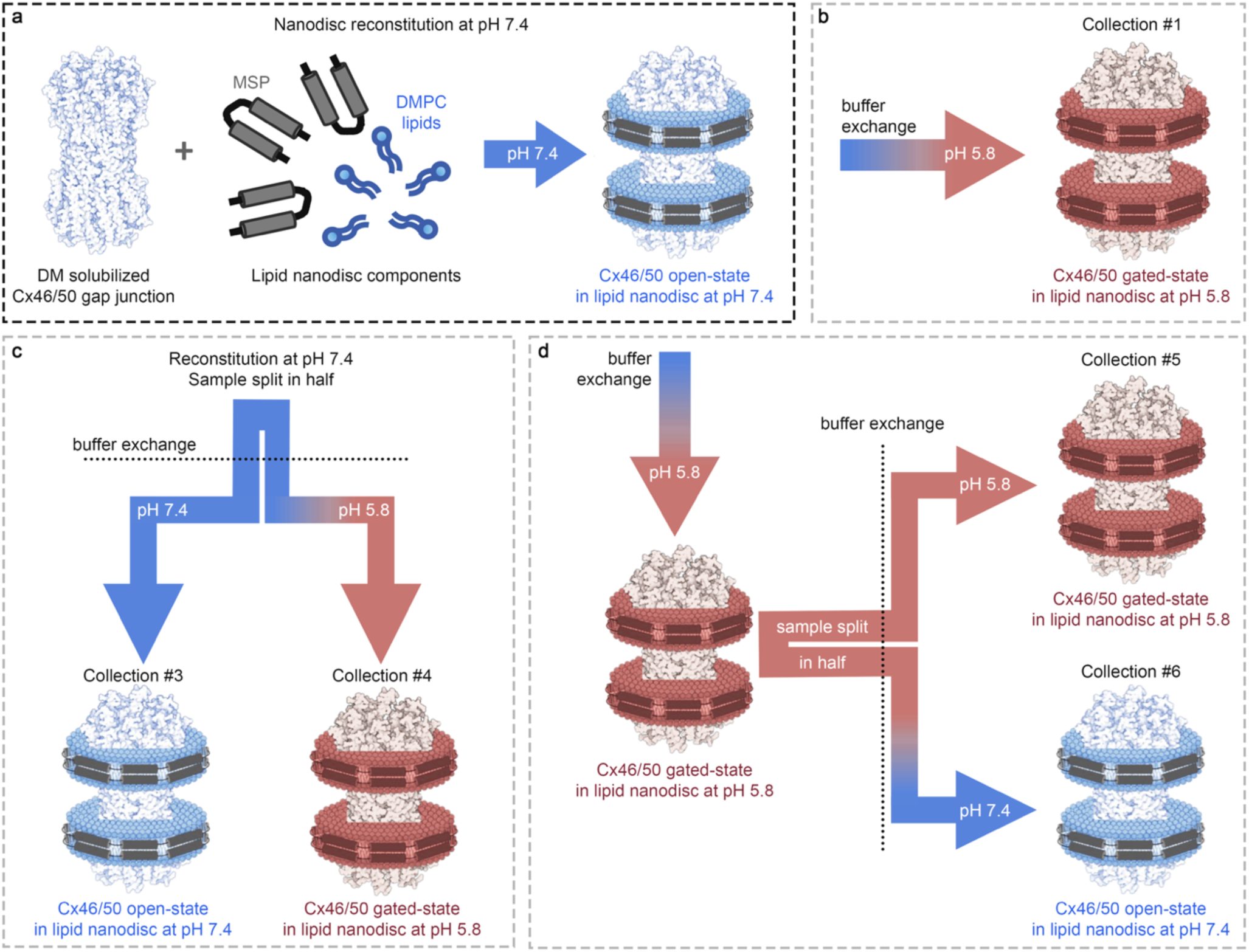
Schematic of experimental design for pH- and lipid-dependent structural studies. **a** Native Cx46/50 gap junction channels were reconstituted into MSP1E1 nanodiscs containing DMPC lipids at pH 7.4. **b** For Collection #1, nanodisc-embedded Cx46/50 channels at pH 7.4 (panel a) were buffer exchanged by gel filtration to pH 5.8 and imaged by cryo-EM. **c** After reconstitution into nanodiscs at pH 7.4 (panel a), the sample was split in half and buffer exchanged by gel filtration to pH 7.4 (control, Collection #3) or pH 5.8 (Collection #4), followed by cryo-EM imaging. **d** Nanodisc-embedded Cx46/50 channels reconstituted at pH 7.4 were exchanged to pH 5.8 by gel filtration to induce gating. The sample was then split: one half was dialyzed into the same pH 5.8 buffer (control, Collection #5), while the other half was returned to pH 7.4 by dialysis (Collection #6). Both samples were then imaged by cryo-EM. Not shown: Cx46/50 gap junctions were reconstituted into amphipols at pH 7.4 and subsequently buffer exchanged to pH 5.8 by gel filtration (Collection #2).

**Extended Data Figure 3.**
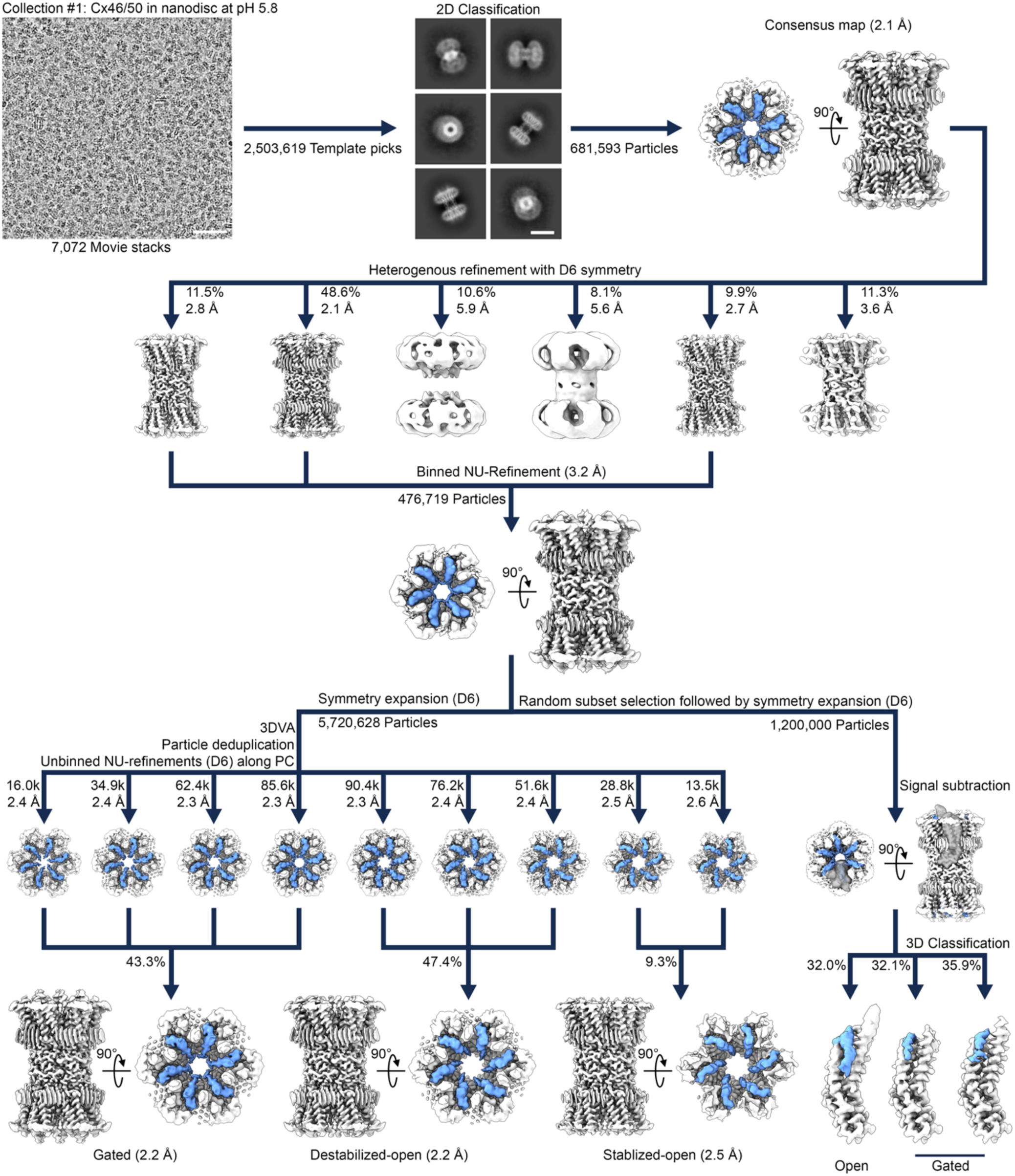
Cryo-EM image processing workflow for the low pH nanodisc dataset (Collection #1). Workflow for the low pH in nanodiscs containing DMPC lipids dataset (Collection #1). Representative cryo-EM micrograph (physical pixel size 0.802 Å^2^; total dose ∼50 e^-^/Å^2^) and 2D classes are shown with 50 nm and 10 nm scale bars, respectively. The dataset was processed using CryoSPARC, with representative maps shown at various stages of the pipeline. After blob picking and template generation (not shown), particles were selected using template picking followed by iterative 2D classification and particle deduplication. Particles underwent successive non-uniform (NU) refinements (D6 symmetry) and unbinning to yield a high-resolution consensus map. Heterogenous refinement (D6 symmetry) was used to isolate high-resolution (< 3.0 Å) particles, which were subsequently binned (1.5 Å pixel size) and subjected to NU-refinement (D6 symmetry). To assess particle heterogeneity at the subunit-level, signal subtraction was performed on a random subset of symmetry expanded (D6) particles, yielding 1.2M monomer particles for signal subtraction and 3D classification. 3D classification revealed primarily gated-state monomers, with a minor population of open-state monomers. To assess particle heterogeneity at the gap junction-level, particles were symmetry expanded (D6) and subjected to 3D variability analysis (3DVA). A mask generated to exclude the nanodisc was used. The primary principal component was divided into nine non-overlapping frames, each locally refined using D6 symmetry. Particles were then deduplicated and subjected to NU-refinement (D6 symmetry). Particles yielding similar 3D reconstructions (classified as gated, destabilized open, and stabilized open-states based on conformation of the NT domain – blue) were pooled and subjected to a final round of NU-refinement with applied D6 symmetry.

**Extended Data Figure 4.**
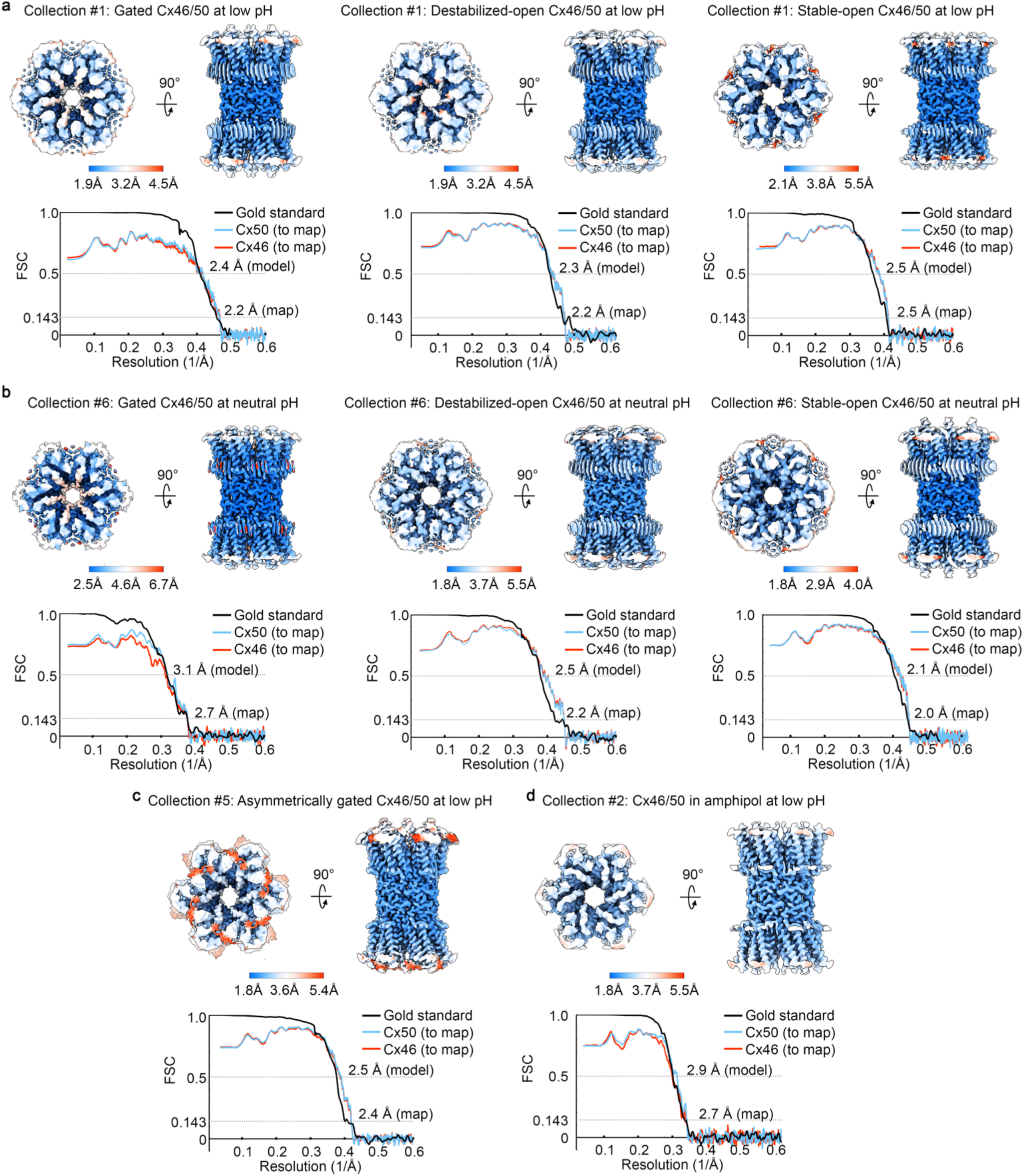
Resolution assessment of cryo-EM 3D reconstructions. Local resolution analysis of final cryo-EM 3D reconstructions using CryoSPARC, displayed by colored surface. Fourier Shell Correlation (FSC) analysis obtained from final 3D reconstructions. Gold-Standard FSC (black) indicating global resolutions (0.143 cut-off). FSC curves comparing atomic models for Cx46 (red) and Cx50 (blue) fit to the respective final cryo-EM 3D reconstruction (0.5 cut-off). Resolution assessment is shown for: **a** (Collection #1) *Left,* gated-state; *middle,* destabilized open-state; *right,* stabilized open-state obtained at low pH. **b** (Collection #6) *Left,* gated-state; *middle,* destabilized open-state; *right,* stabilized open-state obtained at neutral pH. **c** (Collection #5) Representative asymmetrically gated-state obtained at low pH. **d** (Collection #2) Stabilized open-state obtained at low pH in amphipol.

**Extended Data Figure 5.**
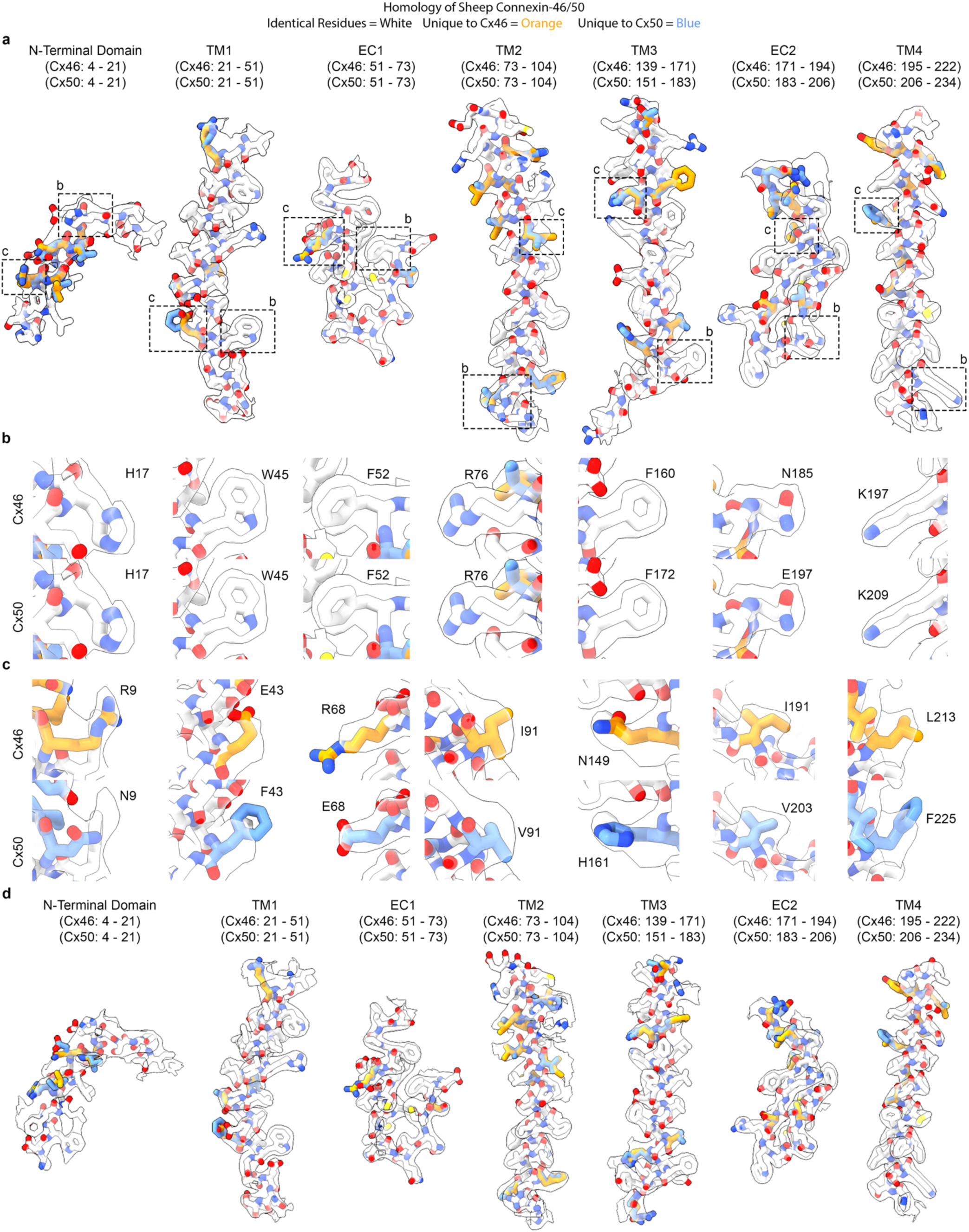
Fit of atomic models to the cryo-EM density map of the low pH gated-state. **a** Segmented cryo-EM map for the low pH gated-state with atomic models for sheep connexin-46 (Cx46) and connexin-50 (Cx50) fit to the experimental densities derived from Collection #1 (2.2 Å, D6-symmetry), including regions for the N-terminal domain, transmembrane domains 1-4 (TM1-4) and extracellular domains 1-2 (EC1-2). Cx46 and Cx50 models are colored according to their pair-wise sequence homology, as being identical (white), or unique (orange for Cx46; blue for Cx50). **b,c** Windows show zoom-views corresponding to boxed regions of the segmented maps, highlighting representative sidechain densities and fit to the atomic models. Regions of identical amino acids (panel b) are fit equally well by both models, whereas regions where the sequence of Cx46 and Cx50 differ (panel c), sidechain density is typically weaker or show features due to imposed averaging of two different sidechains in these areas (or relative flexibility of solvent exposed sidechains). **d** Segmented cryo-EM map for the low pH destabilized open-state with atomic models for sheep Cx46 and Cx50 fit to the experimental densities derived from Collection #1 (2.2 Å, D6-symmetry), including regions for the N-terminal domain, TM1-4, and EC1-2, colored as in panel a.

**Extended Data Figure 6.**
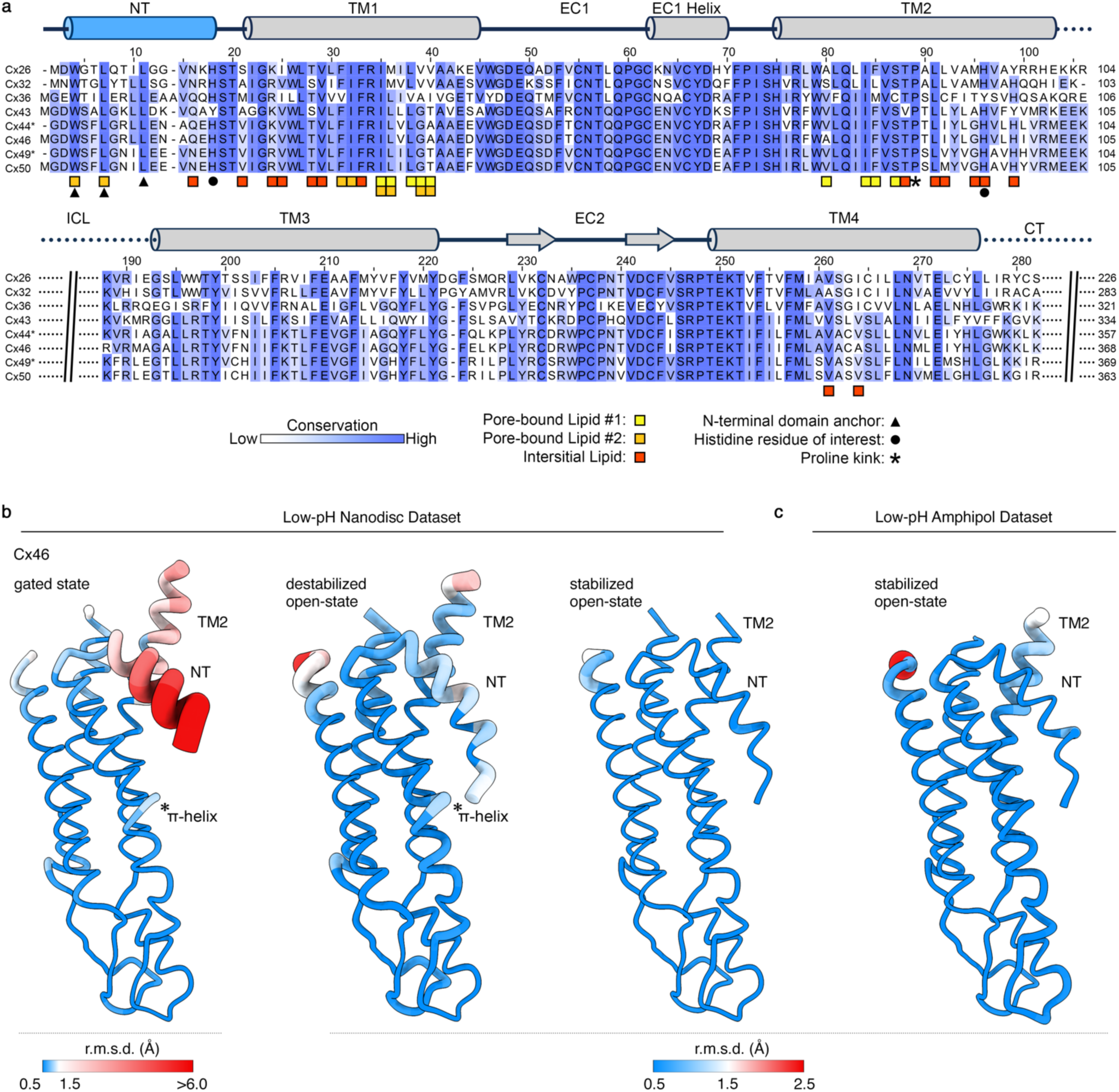
Annotated sequence and structural alignments. **a** Sequence alignment and secondary structure topology of representative human and ovine (Cx44 and Cx49) connexin isoforms, colored by conservation. Conserved hydrophobic residues involved in N-terminal domain anchoring of the stabilized open-state of Cx46/50 are marked with triangles. Conserved histidine residues highlighted in this study (H17 and H95, Cx46/50 numbering) are indicated with circles. The conserved proline kink on TM2 is marked with an asterisk. Residues interacting with pore-bound lipids (PL1 and PL2) are annotated with yellow and orange squares, respectively. Residues interacting with the interstitial lipid (IL) are marked with red squares. **b** Structural alignment and Cα root mean square deviation (r.m.s.d.) analysis of Cx46 in a lipid nanodisc environment obtained in the low pH gated (left) destabilized open (center) and stabilized open (right) states versus the stabilized open state obtained at neutral pH (7JKC). **c** Structural alignment and Cα r.m.s.d. analysis of Cx46 in an amphipol environment obtained at low pH versus the stabilized open state obtained at neutral pH (7JKC). For panels b and c, r.m.s.d. values are visualized by color gradient (blue to red, indicating low to high values) and by the thickness of the backbone in ‘worm’ representation produced in ChimeraX. NT and TM2 domains are labeled. A key for r.m.s.d. values for each set of comparisons is displayed. Asterisk indicates region of minor variability around a π-helix that forms a kink in TM1 (residues ∼39-41) observed in the gated and destabilized open-states.

**Extended Data Figure 7.**
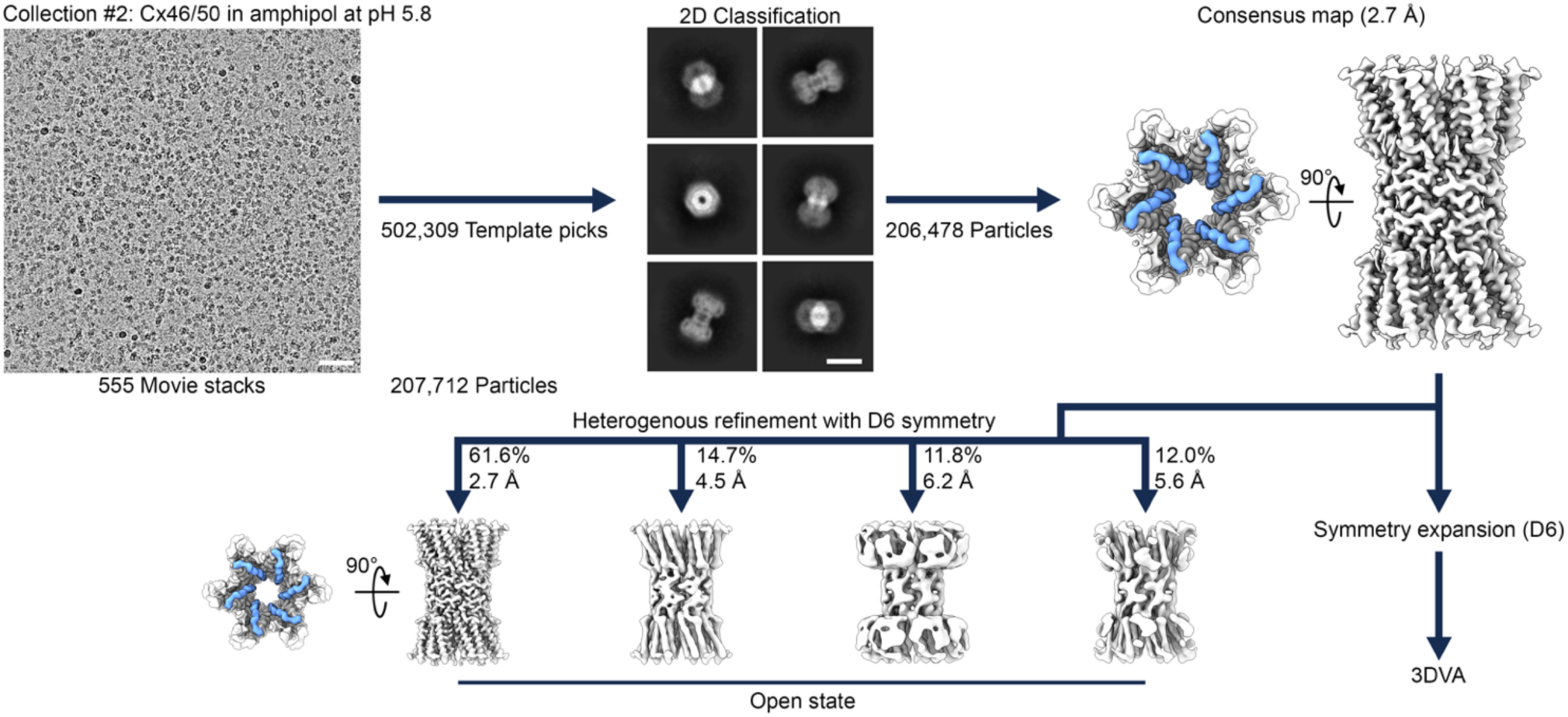
Cryo-EM image processing workflow for the low pH amphipol dataset (Collection #2). Workflow for the low pH in amphipol dataset (Collection #2). Representative cryo-EM micrograph (physical pixel size 0.665 Å^2^; total dose ∼80 e^-^/Å^2^) and 2D classes are shown with 50 nm and 10 nm scale bars, respectively. The dataset was processed using CryoSPARC, with representative maps shown at various stages of the pipeline. After blob picking and template generation (not shown), particles were selected using template picking followed by iterative 2D classification and particle deduplication. Particles underwent successive non-uniform (NU) refinements (D6 symmetry) and unbinning to yield a high-resolution consensus map. Heterogenous refinement (D6 symmetry) was used to isolate high-resolution (< 3.0 Å) particles. This resulted in one class representing the stable-open state. To assess particle heterogeneity at the gap junction-level, particles from the consensus 3D reconstruction were symmetry expanded (D6) and subjected to 3D variability analysis (3DVA). A mask generated to exclude the detergent micelle was used.

**Extended Data Figure 8.**
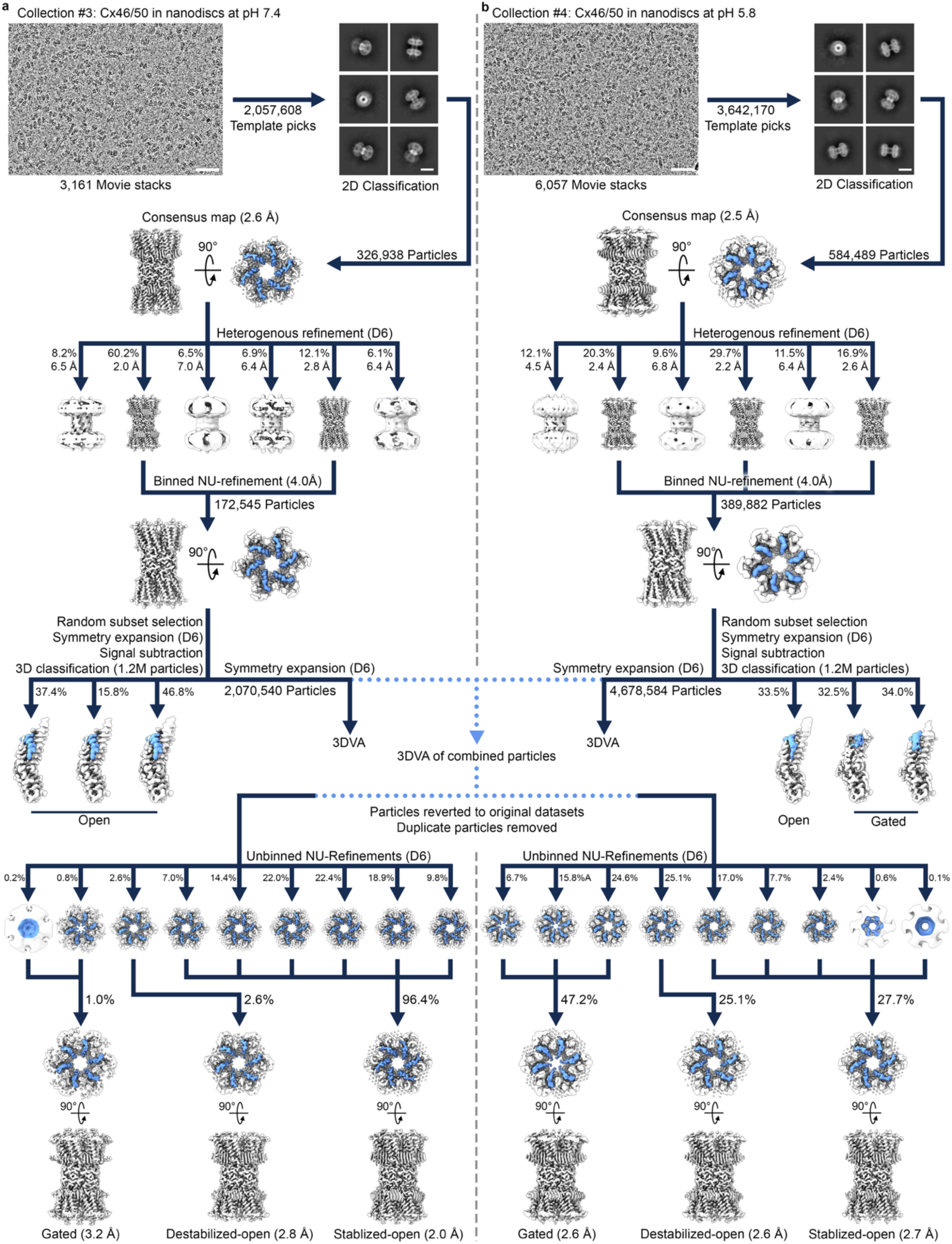
Cryo-EM image processing workflows for the split neutral (Collection #3) and low pH (Collection #4) datasets. **a** Workflow for the neutral pH dataset (Collection #3). Representative cryo-EM micrograph (physical pixel size 0.788 Å²; total dose ∼50 e⁻/Å²) and 2D classes are shown with 50 nm and 10 nm scale bars, respectively. **b** Workflow for the low pH dataset (Collection #4), following the same general procedure as the neutral pH dataset. Representative cryo-EM micrographs and 2D classes are shown at equivalent scale. Both datasets were processed using CryoSPARC, with representative maps shown at various stages of the pipeline. After blob picking, and template generation (not shown), particles were selected using template picking followed by iterative 2D classification and particle deduplication. Particles underwent successive non-uniform (NU) refinements (D6 symmetry) and unbinning to yield a high-resolution consensus map. Heterogenous refinement (D6 symmetry) was used to isolate high-resolution (< 3.0 Å) particles, which were subsequently binned (2.0 Å pixel size) and subjected to NU-refinement (D6 symmetry). To assess particle heterogeneity at the subunit-level, signal subtraction was performed on a random subset of symmetry expanded (D6) particles, yielding 1.2M monomer particles for signal subtraction and 3D classification. For the neutral pH dataset (Collection #3), 3D classification only revealed open-state monomers. For the low pH dataset (Collection #4), 3D classification revealed primarily gated-state monomers, with a minor population of open-state monomers. To assess particle heterogeneity at the gap junction-level, particles were symmetry expanded (D6) and subjected to 3D variability analysis (3DVA). A mask generated to exclude the nanodisc was used for both datasets independently and on the combined datasets. The primary principal component for the combined datasets was divided into nine non-overlapping frames, each locally refined using D6 symmetry. Particles were then reverted to their original datasets, deduplicated, and subjected to NU-refinement (D6 symmetry). Particles yielding similar 3D reconstructions (classified as gated, destabilized open, and stabilized open-states based on conformation of the NT domain – blue) were pooled and subjected to a final round of NU-refinement with applied D6 symmetry.

**Extended Data Figure 9.**
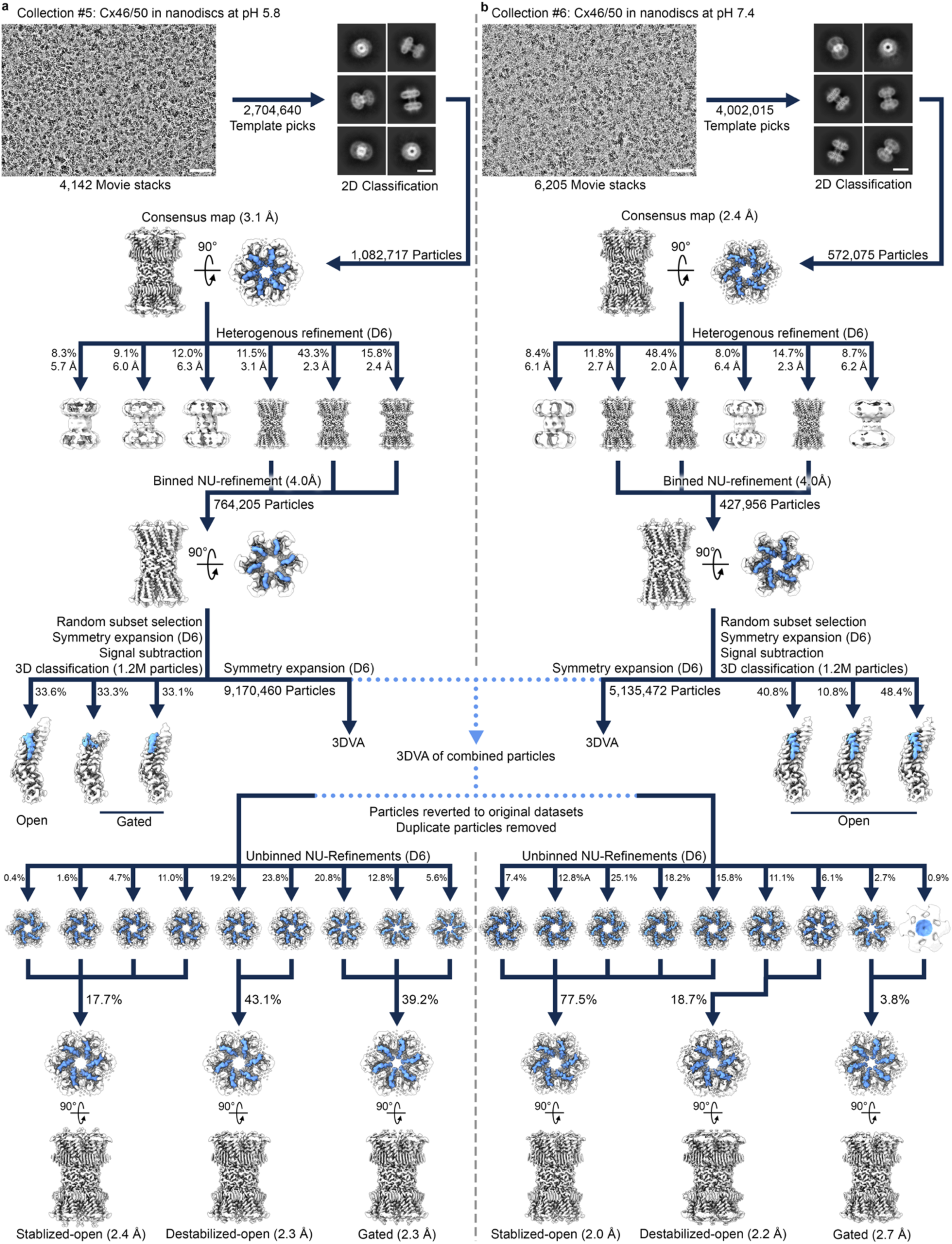
Cryo-EM image processing workflow for the reversibility of lipid-mediated pH gating low pH (Collection #5) and neutral pH (Collection #6) datasets. **a** Workflow for the low pH dataset (Collection #5). Representative cryo-EM micrograph (physical pixel size 0.822 Å^2^; total dose ∼50 e^-^/Å^2^) and 2D classes are shown with 50 nm and 10 nm scale bars, respectively. **b** Workflow for the neutral pH dataset (Collection #6), following the same general workflow as the low pH dataset. Both datasets were processed using CryoSPARC, with representative maps shown at various stages of the pipeline. After blob picking and template generation (not shown), particles were selected using template picking followed by iterative 2D classification and particle deduplication. Particles underwent successive non-uniform (NU) refinements (D6 symmetry) and unbinning to yield a high-resolution consensus map. Heterogenous refinement (D6 symmetry) was used to isolate high-resolution (< 3.0 Å) particles, which were subsequently binned (2.0 Å pixel size) and subjected to NU-refinement (D6 symmetry). To assess particle heterogeneity at the subunit-level, signal subtraction was performed on a random subset of symmetry expanded (D6) particles, yielding 1.2M monomer particles for signal subtraction and 3D classification. For the neutral pH dataset (Collection #6), 3D classification only revealed open-state monomers. For the low pH dataset (Collection #5), 3D classification revealed primarily gated-state monomers, with a minor population of open-state monomers. To assess particle heterogeneity at the gap junction-level, particles were symmetry expanded (D6) and subjected to 3D variability analysis (3DVA). A mask generated to exclude the nanodisc was used for both datasets independently and on the combined datasets. The primary principal component for the combined datasets was divided into nine non-overlapping frames, each locally refined using D6 symmetry. Particles were then reverted to their original datasets, deduplicated, and subjected to NU-refinement (D6 symmetry). Particles yielding similar 3D reconstructions (classified as gated, destabilized open, and stabilized open-states based on conformation of the NT domain – blue) were pooled and subjected to a final round of NU-refinement with applied D6 symmetry.

## SUPPLEMENTAL MOVIE LEGENDS

○ Supplemental Movie 1: 3D variability analysis of Cx46/50 under varying pH and lipid conditions.
○ Supplemental Movie 2: Morph of Cx46/50 models illustrating the lipid-mediated pH gating mechanism.

